# Isolation and characterization of extracellular vesicles from *Caenorhabditis elegans* for multi-omic analysis

**DOI:** 10.1101/476226

**Authors:** Joshua C. Russell, Gennifer E. Merrihew, Julia E. Robbins, Nadia Postupna, Tyek-Kyun Kim, Alexandra Golubeva, Ayush Noori, Kai Wang, C. Dirk Keene, Michael J. MacCoss, Matt Kaeberlein

## Abstract

Cells from bacteria to human release vesicles into their extracellular environment. These extracellular vesicles (EVs) contain multiple classes of molecules, including nucleic acids, proteins, and lipids. The isolation and analysis of EV cargos from mammalian cell culture and liquid biopsy samples has become a powerful approach for uncovering the messages that are packaged into these organelles. However, this approach has not been tenable in invertebrate model systems due to lack of sufficient amounts of pure EVs. Here we report a robust and reproducible procedure to isolate EVs from *Caenorhabditis elegans* with yields similar to those obtained from human cell culture. Through nanoparticle tracking, transmission electron microscopy, flow cytometry, mass spectrometry, RNAseq, and immunoaffinity analysis we provide the first ever detailed characterization of *C. elegans* EV composition and demonstrate that *C. elegans* EVs share fundamentally similar properties with their mammalian counterparts. These include vesicle size, enrichment for lipid rafts, and similar types of RNA and protein cargos. This ability of isolate pure EVs on ascale amenable to multiple types of downstream analyses permits, multi-omics characterization of EV cargos in an invertebrate model system.

## BACKGROUND

The cellular secretion of small membrane-bound extracellular vesicles (EVs) into the external environment is an ancient capacity conserved throughout evolution ^(Deatherage and Cookson, 2012; Schorey et al., 2015),(Robinson et al., 2016)^. EVs range in size from 30-1000 nm in diameter and can be internalized into recipient cells via endocytosis or membrane fusion. There is growing recognition that EVs may play important roles in facilitating intercellular communication through transferring protein, lipid, and genetic cargos ^(Maas et al., 2017),(Mulcahy et al., 2014)^. The content of EVs are influenced by the physiological state of the cells and are thought to play critical roles in diverse cellular processes as well as multiple types of pathological conditions including cancer, immunity, and neurodegenerative diseases.

Many studies have characterized the composition of mammalian EVs. Such EVs are highly enriched in the lipid raft species cholesterol, and sphingomyelin, and in proteins that associate with lipid rafts, including glycosylphosphatidylinositol-anchored (GPI) proteins ^(Wubbolts et al., 2003),(Zhuang et al., 2005),(del Conde et al., 2005)^. The two main types of EVs studied so far are exosomes, which release from the endosomal network and microvesicles which bud directly from the plasma membrane ^(Cocucci and Meldolesi, 2015; van Niel et al., 2018)^. Mammalian exosomes commonly contain membrane proteins such as CD9, CD63, and CD81, as well as lysosomal and endosomal-marking proteins, and various amounts of extracellular matrix proteins while are largely free of nuclear proteins ^(Kowal et al., 2016)^. Microvesicles have less defined protein markers but may contain proteins of mitochondria and endoplasmic reticulum origin ^(Kowal et al., 2016)^. However, the methods utilized to purify EVs do not separate these types of vesicles, so it is currently unclear how to definitively distinguish the cargos from different types of EVs ^(Théry et al., 1999)^. EVs from diverse species, including humans, are known to carry RNA cargos including protein coding mRNAs, and non-coding RNAs including miRNA, rRNA, snoRNA, piRNA etc ^(Zaborowski et al., 2015)(Bayer-Santos et al., 2014)(Figliolini et al., 2014)^. These protein and genetic cargos have been viewed as a rich source of biomarkers because they are thought to reflect the physiological state of their cell of origin.

The nematode *Caenorhabditis elegans* has also been shown to secrete EVs into the external environment ^(Wang et al., 2014),(Buck et al., 2014)^ (Figure 1A). For example, EVs secreted from ciliated sensory neurons contain signals that influence the behavior of male nematodes ^(Wang et al., 2014)^. In *C. elegans* microvesicles are shed from the plasma membrane in a flippase-dependent manner ^(Beer et al., 2018),(Wehman et al., 2011)^ while exosomes deliver signaling factors that are necessary for proper cuticle development ^(Liégeois et al., 2006)^. A thorough review of EV-signaling in invertebrate model systems has been recently published ^(Beer and Wehman, 2017)^. These investigations have begun to elucidate the genetic pathways and physiological impacts of microvesicle and exosome signaling. However, the protein, lipid, and genetic cargos of invertebrate EVs remains largely unknown due to a lack of methodology for obtaining sufficient amounts of pure EVs. Here we report the first large-scale purification and multi-omics characterization of *C. elegans* vesicles and provide evidence for highly conserved features of EV structure and function.

**Figure 1:**
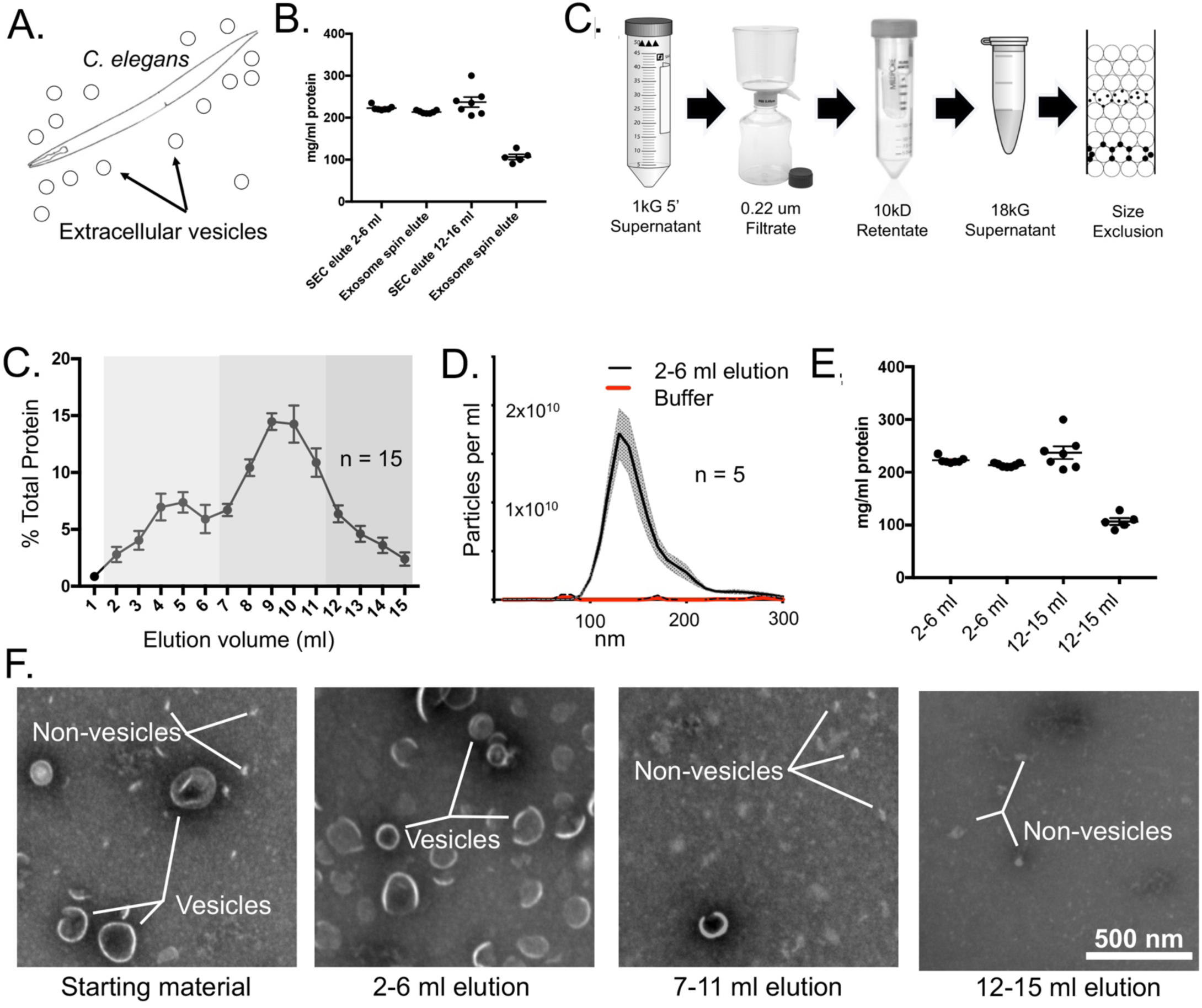
Purification of vesicle-like particles from C. *elegans* secretate. A) C. *elegans* secretes extracellular vesicles into their outside environment. B) Fractionation scheme to enrich for secretate particles. C) Protein elution profile from 10 ml Sepharose CL2-Bcolumn loaded with 1ml of secretate. The fractions that are consolidated in further analyses are shaded into groups D) Nanoparticle tracking analysis of the consolidated Sepharose elution fractions 2-6 ml. 95% confidence interval is shaded gray E) The consolidated Sepharose elution 2-6 ml fractions were then passed over a commercial exosome spin column. The total protein contained in material run over the exosome spin column and the material elution from the spin column are not significantly different (n = 12 biological replicates) Error bars S.E.M. F) TEM analysis of the consolidated Sepharose elution fractions. The starting material has both vesicle-like particles and non-vesicles particles. The consolidated 2-6 ml elution fraction were enriched for vesicles with little non-vesicle particles. The 7-11 ml consolidated fractions contained few vesicle-like particles but many non-vesicle particles. The 12-15 ml elution fractions contained only non-vesicle particles. All micrographs shown at the same magnification. Scale bar 500nm.

## RESULTS

### Isolating EVs from *C. elegans*

In order to generate sufficient biomass for purifying EVs, we cultivated large populations (~500,000) of age-synchronized worms on high-growth plates seeded with NA22 bacteria at 20 C until they reached young adulthood. We then washed the worms off the plates and removed bacteria and other debris through sucrose floatation ^(Zanin et al., 2011)^. The number of worms was quantified and then placed on a rotator in S Basal with 5 μg/mL cholesterol at a density of 1 worm per microliter. We allowed the worms to secrete EVs into the S Basal media for 24 hours after which time the worms were pelleted at 2000 X G for 5 minutes and assessed for viability. Nearly all of the animals were uniformly vigorous when transferred to NGM plates. We quantified the viability of these transferred animals by determining how many did not move from the spot deposited within five minutes. In three separate biological replicates with over 200 animals we did not observe one that did not move from the deposit site, indicating that the 48-hour incubation did not damage the animals.

### FRACTIONATION

We chose to focus on the small (<200 nm) EVs because of their reported physiologic roles ^(Hyenne et al., 2015),(Liégeois et al., 2006),(Wang et al., 2014)^. Therefore, we filtered the supernatant through a 0.22 μm filter to remove larger particles. Recently it was reported that ultrafiltration, rather than ultracentrifugation, results in less damage to vesicles and a higher recovery rate ^(Benedikter et al., 2017)^. While most ultrafiltration substrates will bind EVs, regenerated cellulose does not ^(Vergauwen et al., 2017)^, so we decanted and concentrated the 0.22 μm filtrate using a 10 kD mwco regenerated cellulose filter to the final volume of 1 ml at 4C (MilliporeSigma, Burlington MA, USA; Cat # UFC901008) (Figure 1B). To prevent protein degradation, we then added HALT protease inhibitor cocktail and EDTA (Thermo Fisher Rockford IL, USA Cat # 78430).

### Size Exclusion Chromatography

To separate any EV-sized particles from soluble proteins or lipids we replicated a method that was developed for isolating EVs from human blood ^(Böing et al., 2014)^. We reasoned that by replicating these experimental conditions any small vesicles in our concentrated secretate should elute at a similar volume. We passed the retentate over a 10 mL Sepharose CL-2B size-exclusion column and used ice-cold S Basal with HALT protease inhibitor cocktail and EDTA as the mobile phase and collected 15 × 1 mL fractions (MilliporeSigma, Burlington MA, USA; Cat # CL2B300-100ML). Qubit analysis on each 1 mL fraction showed that our elution profile was similar to human EVs ^(Böing et al., 2014)^, with a small peak around 4 mL and a large broad peak encompassing 8-15 mls (Figure 1C). sized lipoprotein aggregates, we examined them by transmission electron microscopy (TEM). When vesicles are prepared for TEM they take on a characteristic cup shape due to dehydration, while solid particles appear as bright punctate dots with negative stain ^(Mathivanan et al., 2010),(Kalra et al., 2013)^. To analyze our size exclusion column elution, we spotted 2 μL of the fractions onto glow discharged formvar-carbon coated copper mesh grids (Polysciences, Warrington, PA USA; Cat # 24915-25), stained with 2% phosphotungstic acid (PTA) adjusted to pH 7.0 (Ted Pella Redding, CA USA; Cat # 19402) and washed the grids three times with 2 μL filtered MilliQ water. We imaged our samples at 19,000 X with a Philips CM100 TEM and found that the elution fractions from 2 mL to 6 mL contained abundant ~100 nm spherical cup-shaped particles while the later fractions did not (Figure 1F). The results showed that the size exclusion column effectively separated the cup-like EVs from these non-vesicle particles. It is likely that these non-vesicle structures are high density lipids because this column set-up has been shown to concentrate EVs in the 2-6 mL elution fractions and concentrate high-density lipids in the 7-12 mL fractions ^(Böing et al., 2014)^. Representative images from three biological replicates is presented in Supplemental (Supplemental Figure 1).

### Nanoparticle tracking analysis

To determine the number and size distribution of the particles eluted from the size exclusion column, we performed nanoparticle tracking analysis (NTA). Our TEM results suggested that the particles were confined to elution volumes between 2-6 mLs. Therefore, we consolidated the 2-6 mL elution fractions and concentrated to 1 mL using a nitrocellulose spin column (MilliporeSigma, Burlington, MA; Cat # UFC801024). In order to get the particle concentration within the working range of the NanoSight ns3000 we diluted our samples 1:100 with MilliQ water just prior to analysis. We examined five biological replicates and found that they all had one main monodisperse population of particles with a mean size of 150 nm (Figure 1D). Coupled with our previous TEM results, this suggests our EV purification method effectively isolated lipid vesicles from *C. elegans* that have the characteristic size and morphology of exosomes and small microvesicles.

### Vesicle fractions are highly enriched for EV protein

Although our NTA and TEM analysis suggested that we had greatly enriched for particles we wanted to determine whether there were still freely-soluble proteins in our particle samples. To determine this, we consolidated the 2-6 mL elution fractions, concentrated the volume to 100 μL over a 10kD regenerated nitrocellulose membrane (MilliporeSigma, Burlington MA, USA; Cat # UFC801024), and then passed them over a small (100 μL sample volume) commercial exosome size exclusion spin column that has been shown to separate EV sized particles from freely soluble molecules (Invitrogen, Carlsbad CA; Cat # 4484449) ^(Roberts-Dalton et al., 2017),(Kenigsberg et al., 2017)^. We found that the total protein eluted from the spin columns was not statistically different from the amount of total protein loaded, suggesting that the protein contained in the consolidated 2-6 mL elution fractions is almost entirely associated with exosome-sized particles while over half of the protein from the 12-16 mL fraction was retained on the column (Figure 1E).

### Flow cytometry analysis reveals that purified particles are detergent-soluble

Due to lack of prior flow cytometry studies on *C. elegans* EVs, we first validated our methodologies with human cell culture EVs. We conducted our human cell line EV-isolation scheme as described above on conditioned media from human iPS-derived neuronal cell cultures as starting material. Samples were analyzed using an Apogee A50 flow cytometer (Apogee Flow Systems, Northwood UK). The Apogee A50 is designed for resolving EV-sized nanoparticles and has been routinely utilized for characterizing EVs ^(Chandler et al., 2011),(Dabrowska et al., 2018).(Surman et al., 2018)^. Flow cytometer sheath solutions were 0.1 um filtered before use. Polystyrene fluorescence beads (0.50 um) and size calibrated non-fluorescent silica beads (0.18, 0.24, 0.30, 0.59, 0.88, and 1.30 μm) were resolved with small angle light scattering (SALS) and large angle light scatter (LALS) (Apogee Hempstead UK; Cat # Cat #1493). Because vesicles have a different refractive index than the beads, they resolve differently with light scattering. Therefore, we did not use the calibration beads as a means to determine absolute size of particles but simply as a way to check the performance and consistency of the Apogee A50 before analyzing our experimental samples, running the calibration beads prior to each day’s experiment. We conducted all flow cytometry experiments with the same flow rate (1.5 μL /min) and accumulation time (180 seconds) so that we could compare the relative concentrations of particles between different runs.

First, we examined the small angle and large angle light scattering of consolidated 2-6 mL elution fractions from conditioned cell culture media. This revealed a single highly enriched particle population comprising over 80% of total particle events (Supplemental Figure 2). We then incubated the samples with the lipophilic fluorophore DI-8-ANEPPS that was recently shown to quantitatively label EVs(de Rond et al., 2018) (Thermo Fisher Rockford IL, USA; Cat # D3167). The dye was added to a final concentration of 500 nM. We found that over 80% of the total particle events were strongly fluorescent (Supplemental Figure 2). To confirm that these fluorescent events represent lipid vesicles and not similarly-sized solid lipid particles, we treated Di-8-ANEPPS-stained particles with 0.05% v/v Triton X-100. This reduced dye-labeled particle counts to background levels, verifying that our purification scheme results in abundant, EVs from cultured cells (Supplemental Figure 2).

**Figure 2:**
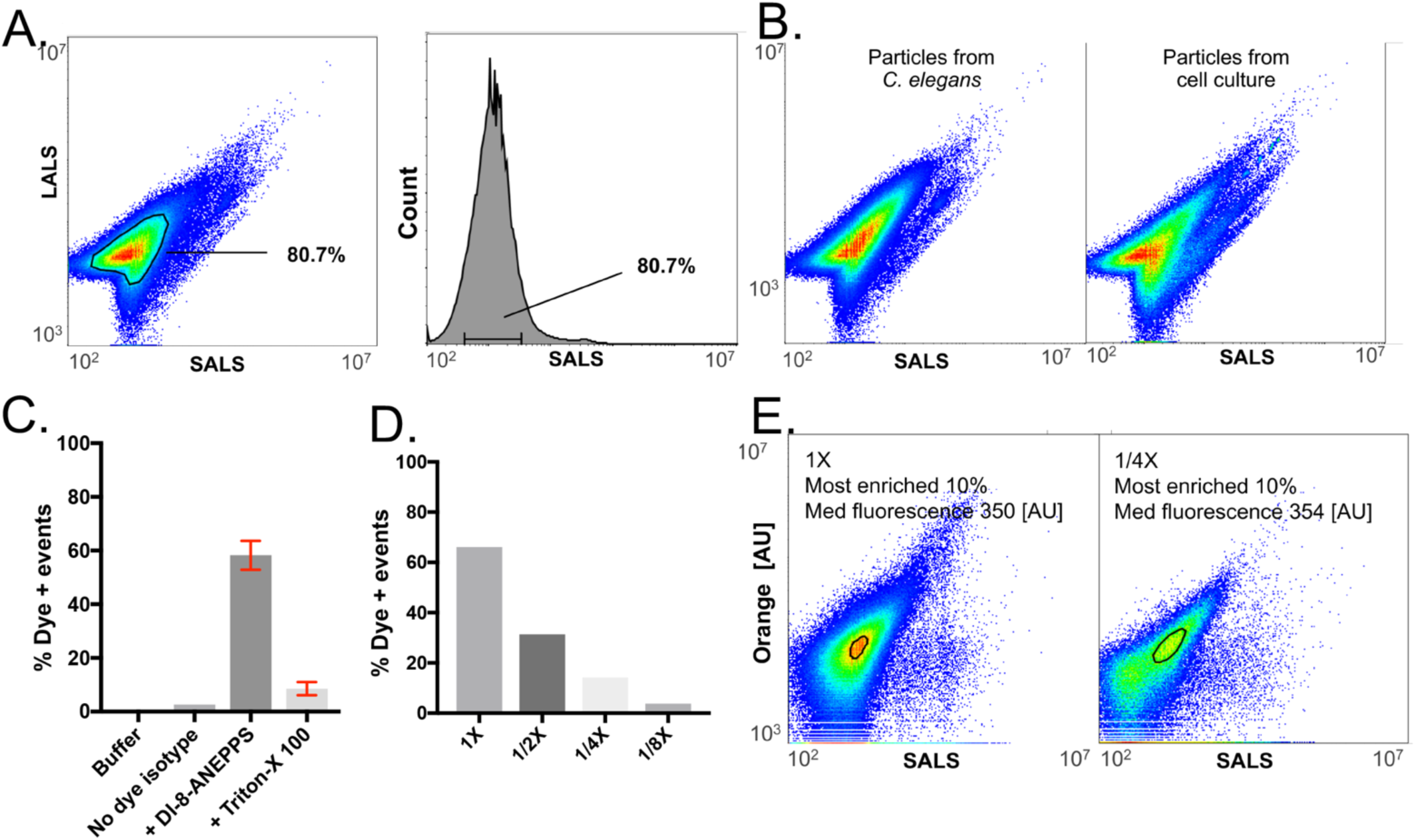
Flow cytometry analysis reveals secretate particles purified from *C. elegans* secretate are vesicles. A) Small and large-angle scattering reveals a single population of particles in the *C. elegans* secretate. Over 80% of the particles are contained within the most-dense population. Light scattering and histogram of particle events separated by SALS A) Particles purified from C. elegans secretate or human cell culture conditioned media display matching light scattering properties. C) DI-8-ANEPPS robustly labels C. *elegans* EVs. Gating on DI-8-ANEPPS fluorescence reveals that the purified particles from C. *elegans* secretate are robustly labeled with fluorescence and that these events are reduced to background with the addition of 0.05 % Triton-X 100 and light sonication. D) The events decreased in accordance with dilution factor. The mean intensity and the SALS of individual particle events from the hlghest-denslty population hot-spot (comprising almost enriched 10% population of particle events) did not change. This indicates that we can resolve single extracellular vesicles with FACS.

We next sought to determine if comparable FACS methods could be used to analyze our isolated *C. elegans* secretate. Our TEM and NTA results indicated the particles purified from the *C. elegans* secretate are comparably-sized to small EVs from human cell culture media. Given the structural simplicity of EVs (no internal membrane structures) we predicted that the purified particles from *C. elegans* should distribute similarly across the size-angle and low-angle light scattering axes as our human iPSC-derived EVs. Indeed, small and large angle light scattering revealed that, like hiPSC-derived EVs, over 80% of the *C. elegans* particles were concentrated into a dense population. The distribution of these events across the small angle light scattering revealed a sharp monotonic distribution similar to the size distribution identified in our NTA experiments. The small and long light scattering distributions of *C. elegans* particles were almost identical to hiPSC-derived EVs indicating they have similar sizes and refractile properties (Figure 2A, Supplemental Figure 3).

**Figure 3:**
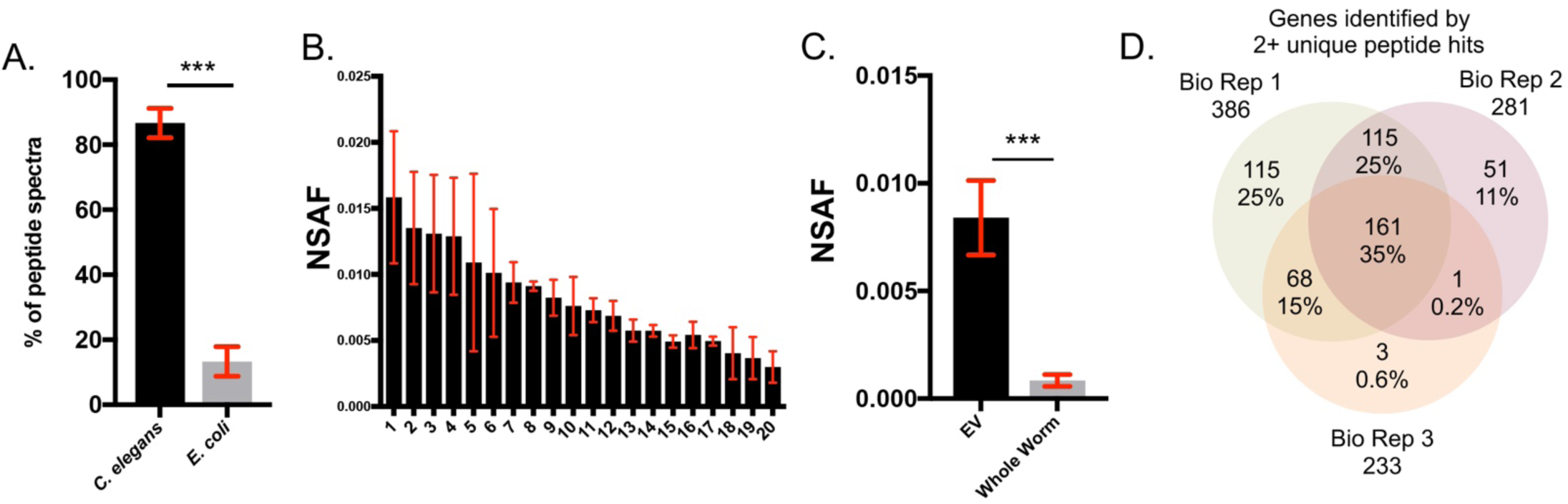
Proteomics analysis *of C. elegans* **EVs.** A) The relative levels of the proteins are similar between runs. Number of genes identified in EV proteomic experiments by two or more unique peptide hits B) Absolute NSAF scores of the 20 most-abundant genes across three biological replicates reveals. Error bars standard deviation. C) Comparison of NSAF value rations between EVs and whole worm lysate D) Venn diagram of the intersection of genes identified in biological replicates. 35% of genes were found in all three replicates

We then wanted to determine if DI-8-ANEPPS could also be used as a general EV marker for *C. elegans*. We exposed the isolated particles to DI-8-ANEPPS and analyzed under the same conditions as our hiPSC-derived EVs FACS experiments. We found that DI-8-ANEPPS labeled particles robustly, highlighting over 60% of total particles. These particles were disrupted by 0.05% Triton-X 100 and light sonication indicating that the particles isolated from the *C. elegans* secretate are almost entirely EVs (Figure 2B). To test whether we were measuring single particles or coincident events, a confounding factor when analyzing nanoparticles with FACS ^(Van Der Pol et al., 2012)^, we performed a series of two-fold dilutions of DI-8-ANEPPS-stained EVs. The measured event rate decreased in proportion to the dilution (Figure 2C); however, the median fluorescence of the most-dense 10% population of DI-8-ANEPPS-labeled particles did not significantly change. This demonstrates that successful quantification of single EVs.

### Proteomic analysis of *C. elegans* EV cargos

To determine the spectrum of protein cargos in *C. elegans* EVs, we conducted proteomic analysis on isolated EVs from three biological replicate cohorts of worms. Because our samples contained protease inhibitors, which can interfere with Trypsin digestion for mass spectrometry analysis, we separated proteins from the inhibitors through SDS-PAGE. The protease inhibitors have a very low molecular weight and so run well below the protein bands. Purified EV fractions were extracted in RIPA buffer (MilliporeSigma; Cat # 20-188) and NuPage LDS 4X sample buffer (ThermoFisher; Cat # NP0008) and reducing agent (ThermoFisher; Cat # NP0009) before heating to 70C for 10 minutes. Samples were spun at 18kG for 15 minutes at 4C. Supernatant was removed and loaded onto 10% NuPage SDS-PAGE (ThermoFisher; Cat # NP0301PK2) with large sample capacity. The samples were run 1 cm into the gel. The gel was then washed twice for 60 min in 250 mL of MilliQ water on an orbital shaker. Each lane was excised and then trypsin digested (Promega, Madison, WI; Cat # V5280). SDS was removed with SDS removal columns (Pierce, Rockville, Il, USA; Cat # 87777) and salts were removed with MCX columns (Waters, Milford, MA, USA; Cat # 186002051). The peptides from each fraction were analyzed using a 30 cm fused silica 75 μm column and a 4 cm fused silica Kasil1 (PQ Corporation, Malvern, PA, USA) frit trap loaded with Repro sil-pur C18 reverse phase resin (Dr. Maisch, Gmbh, Germany) with a 120-min LC-MS/MS run on a Thermo LTQ-Orbitrap Velos mass spectrometer coupled with a Waters nanoACQUITY UPLC system. The MS/MS data was searched using COMET against a FATSA database from WormBase plus contaminate proteins. P-values and q-values were assigned to PSMs and peptides with a 1% FDR using Percolator ^(Käll et al., 2007).^ More than 80% of the peptide spectra corresponded to *C. elegans* peptides, with the remainder mapping to *E. coli* peptides (Figure 3A), indicating that our protocol successfully removed a majority of the bacterial EVs. The complete proteomic data sets are included in Supplemental. The unique *C. elegans* protein hits were then ranked according to their Normalized Spectral Abundance Factor (NSAF). This methodology gives a strong qualitative assessment of the relative protein abundance between biological replicates because it normalizes for the overall peptide abundance in each sample as well as for the size of the proteins ^(McIlwain et al., 2012),(Florens et al., 2006)^. The peptide abundances of the top proteins in the three biological replicates were relatively consistent. Figure 3B depicts the NSAF score for 20 proteins across all three replicates with standard deviation about 40% standard deviation (Figure 3B). We found that the NSAF values in our EV samples were all much greater than those from the whole worm lysate samples, with an average enrichment around 70-fold (Figure 3B; Supplementary Table 1). The strong enrichment of these proteins compared to whole worm lysate suggests that *C. elegans* actively and selectively load protein cargo into EVs. The results of these analyses are publicly available (https://chorusproject.org/anonymous/download/experiment/d41c70cbecfe42a0a00050488903a163)

**Table 1:** GO analysis of *C. elegans* EV proteins. A) EV proteins are enriched for proteolysis, stress, and metabolic processes B) EV proteins are enriched for peptidase, carbohydrate binding, and catalytic activity C)The EV proteins are most associated with the membrane raft, whole membrane, and extracellular cell components D) The KEGG pathways of EV proteins include carbon metabolism, amino acid biosynthesis, and other metabolic pathways E) The EV proteins are enriched for PFAM protein domains that are associated with lectin binding, glycoproteins and proteases.

| #pathway ID | Pathway description | Gene count | False discovery rate |
| --- | --- | --- | --- |
| GO.0006508 | Proteolysis | 39 | 4.52E-20 |
| GO.0006952 | Defense response | 32 | 1.30E-18 |
| GO.0006950 | Response to stress | 42 | 3.25E-17 |
| GO.0044712 | Single-organism catabolic process | 24 | 1.67E-16 |
| GO.0071704 | Organic substance metabolic process | 91 | 6.75E-15 |
| GO.0009056 | Catabolic process | 34 | 1.35E-13 |
| GO.0008152 | Metabolic process | 106 | 7.20E-13 |
| GO.0044238 | Primary metabolic process | 85 | 1.06E-12 |
| GO.0019752 | Carboxylic acid metabolic process | 24 | 1.12E-12 |
| GO.0045087 | Innate immune response | 20 | 3.39E-11 |
| GO.1901575 | Organic substance catabolic process | 30 | 3.39E-11 |
| GO.1901564 | Organonitrogen compound metabolic process | 32 | 3.43E-10 |
| GO.0044724 | Single-organism carbohydrate catabolic process | 10 | 3.62E-10 |
| GO.0005975 | Carbohydrate metabolic process | 21 | 9.02E-10 |

**B) Molecular function**
| #pathway ID | Pathway description | Gene count | False discovery rate |
| --- | --- | --- | --- |
| GO.0008233 | Peptidase activity | 39 | 2.86E-24 |
| GO.0016787 | Hydrolase activity | 62 | 6.00E-20 |
| GO.0030246 | Carbohydrate binding | 30 | 1.16E-15 |
| GO.0003824 | Catalytic activity | 93 | 1.58E-15 |
| GO.0008238 | Exopeptidase activity | 12 | 1.77E-11 |
| GO.0004185 | Serine-type carboxypeptidase activity | 7 | 2.04E-09 |
| GO.0003674 | Molecular function | 136 | 5.41E-09 |

C) Cell component
| #pathway ID | Pathway description | Gene count | False discovery rate |
| --- | --- | --- | --- |
| GO.0045121 | Membrane raft | 19 | 2.96E-23 |
| GO.0098805 | Whole membrane | 20 | 3.08E-09 |
| GO.0005576 | Extracellular region | 21 | 2.43E-07 |
| GO.0005737 | Cytoplasm | 50 | 5.73E-07 |
| GO.0030016 | Myofibril | 10 | 3.09E-06 |

D) KEGG pathway
| #pathway ID | Pathway description | Gene count | False discovery rate |
| --- | --- | --- | --- |
| 1200 | Carbon metabolism | 17 | 2.63E-14 |
| 1230 | Biosynthesis of amino acids | 15 | 2.63E-14 |
| 4142 | Lysosome | 14 | 9.97E-13 |
| 1100 | Metabolic pathways | 32 | 1.23E-10 |
| 10 | Glycolysis / Gluconeogenesis | 9 | 6.70E-09 |
| 630 | Glyoxylate and dicarboxylate metabolism | 7 | 1.76E-07 |

**Table 1: GO analysis of *C. elegans* EV proteins.**
| #pathway ID | Pathway description | Gene count | False discovery rate |
| --- | --- | --- | --- |
| PF00059 | Lectin C-type domain | 24 | 8.47E-18 |
| PF03409 | Transmembrane glycoprotein | 9 | 3.77E-10 |
| PF00026 | Eukaryotic aspartyl protease | 6 | 2.25E-05 |
| PF00112 | Papain family cysteine protease | 6 | 4.54E-05 |
| PF00147 | Fibrinogen beta and gamma chains | 4 | 4.54E-05 |

### Gene ontology and enrichment analysis

These proteomic results were then filtered for proteins identified with two or more unique peptide hits. To convert the protein hits to a set of expressed genes the different protein isoforms were all condensed to their encoding gene. This resulted in an average of 380 genes corresponding to each of the three biological replicates. A total of 161 genes were identified in all three biological replicates constituting 34% of the total uncovered (Figure 3C). Overall 68% of the identified genes were shared between two sets. We analyzed the gene ontology (GO) annotations from the set of 161 high-confidence genes whose proteins were identified in all three biological replicates using string-db.org ^(Szklarczyk et al., 2017)^. The proteins were significantly enriched into functional networks with significantly more interactions than expected by chance (PPI enrichment P-value < 1 × 10^−16^). The “cellular component” most enriched in *C. elegans* EVs is the membrane raft, which is consistent with prior work indicating that mammalian EVs are enriched for lipid rafts ^(Skotland et al., 2017),(Pfrieger and Vitale, 2018)^. Other cellular components included the lysosome, vacuole, whole membrane, and extracellular region (Table 1A). Biological pathway GO analysis revealed numerous metabolic and catabolic processes as well as stress response, defense response, and innate immune response. (Table 1B). The most-enriched KEGG pathway was carbon metabolism, followed by several other metabolic and catabolic pathways (TABLE 1C) Some of the molecular functions most enriched are peptidase, hydrolase, and carbohydrate binding (Table 1D). The most enriched protein domains were the Lectin C-type domain and transmembrane glycoprotein (Table 1E).

We next asked whether *C. elegans* EVs contain orthologs of known human EV proteins. To do this, we determined the reciprocal BLAST best hits of the 100 proteins most identified in human EV studies and uploaded to Exocarta.org. 48 of the 100 proteins had reciprocal best hits. We then compared the proteins hits from our samples and found that 11 (~24%) of them also present in our proteomics experiments (Supplementary Table 3A). Although reciprocal best hits are a stringent measure of orthology this approach can overlook protein families with strongly conserved sequences because none of them are distinguished as the best hit (e.g. actin). Therefore, we then conducted BLAST analysis against each of the full-length human proteins and found that 85 of these had *C. elegans* orthologs with a value less than e^−30^. We then determined how many of these genes were identified in our EV proteomics samples. 31 of the 85 human EV proteins were represented in our proteomics results. A table of the orthologs along with the individual human protein sequences and *C. elegans* blast results is included in supplemental (Supplementary Table 3B). We then used a Fisher’s exact test to determine the significance of the overlaps between human and worm proteins with best reciprocal hits. This returned a P-value 2.36 × 10^−9^ indicating our proteomic peptide matches were significantly enriched for frequently observed proteins in human EV studies. The two-by-two matrix used to calculate the P-value is in supplemental (Supplementary FIG 8).

### Canonical human EV transmembrane proteins can be used to identify *C. elegans* EVs

The tetraspanin CD63 is a canonical marker of EVs in human research. *C. elegans* has 20 tetraspanin genes, however they have not been examined in the context of EV signaling ^(Hemler, 2005)^. Therefore, we conducted Western analysis with antibodies against the canonical human EV-marking tetraspanins anti-CD9, anti-CD63, and anti-CD81 on whole worm lysates. While the anti-CD9 and anti-CD81 showed no immunoreactivity the anti-CD63 showed bands of similar size to previous human EV studies. Therefore, we analyzed purified EVs for anti-CD63 immunoreactivity. The purified EVs showed a strong single band at the expected size (Figure 4C). To determine if this human EV-marking reagent could also identify intact EVs we incubated our EV fraction with commercial Alexa Fluor-conjugated anti-CD63 (BD Biosciences San Jose, CA; Cat# 561983). This method has been used in the past in FACS analysis of human EVs ^(Logozzi et al., 2009),(Tian et al., 2018),(Clayton et al., 2001),(Pospichalova et al., 2015)^. About a third of the total events between 100 and 300 nm were anti-CD63 positive and these positive events disappeared with Triton-X 100 treatment, indicating that this anti-CD63 reagent can also be used to bind *C. elegans* EVs (Figure 4D).

**Figure 4:**
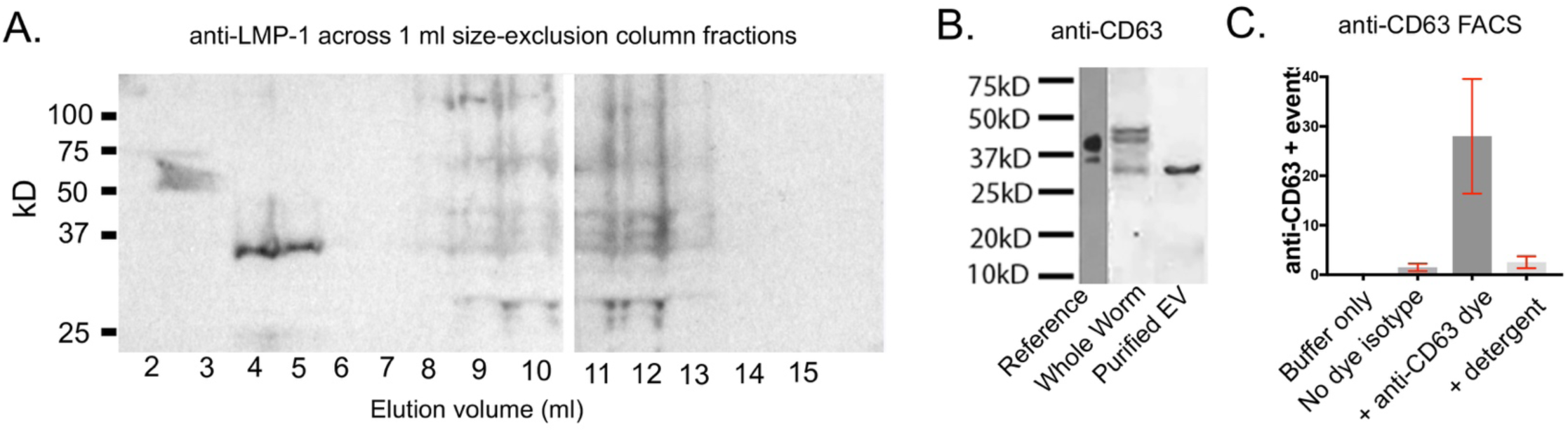
Protein cargos of EVs are similar between humans and *C. elegans*. A) EV-containing SEC fractions (Elution volumes 3 mL and 4 ml) are immunoreactive with for anti-LMP-1 while other later elution fractions are not. B) antl-CD63 Western shows strong immunoreactivity at the expected size. C) Flow cytometry reveals that detergent-sensitive particles are labeled with the human EV membrane marker anti-CD63-Alexa Flour 647

Our mass spectrometry experiments also identified unique LMP-1-derived peptides in all biological replicates. LMP-1 is the homolog of a commonly-identified mammalian exosome membrane protein LAMP-1 ^(Kostich et al., 2000),(Leone et al., 2018),(Vonk et al., 2018)^. To determine whether LMP-1 was specifically associated with EVs or was also in other size exclusion fractions we conducted Western analysis with a monoclonal anti-LMP-1 antibody against all 15 1 mL SEC elution fractions (Developmental Studies Hybidoma Bank, Iowa City, Iowa, Cat # LMP-1). We found strong immunoreactivity only within the elution fractions that are enriched for EVs. This suggests that within the context of *C. elegans* secretate LMP-1 could be an EV-specific membrane-marker (Figure 4A).

### elegans EVs contain RNA

A key feature of mammalian EVs is their ability to convey RNA cargos between cells ^(Raposo and Stoorvogel, 2013)^. To quantify EV-associated RNA we processed our purified samples with Total exosome protein and RNA isolation kit (Invitrogen, Carlsbad CA, USA, Cat # 4478545). We then characterized the abundance and size of the elutes on an Agilent 2200 TapeStation (Agilent, Santa Clara CA, USA; Cat # G2991AA) using high-sensitivity screen tape (Agilent, Santa Clara USA Cat # 5067-5579) along with a small RNA calibration ladder (Agilent, Santa Clara CA, USA; Cat # 5067-1550). The vesicle samples displayed substantial small RNAs as well as 16s and 28s rRNA species at the expected sizes. To determine what RNA species are packaged into EVs we conducted RNAseq. We used a Qiagen small RNA Sample Preparation kit to prepare a cDNA library from our EV-associated RNA (Qiagen Germantown MD, USA Cat # 331502) while an Illumina MiSeq v2 kit (300 cycles) was used to prepare the sequencing library with 5’ adapter sequences (San Diego USA; Cat# MS-102-2002). The sample was then submitted for single-end sequencing on an Illumina MiSeq desktop sequencer. Adapter trimming and sequence alignment was conducted in the sRNAnalyzer ^(Wu et^ al., 2017).

A majority of the reads from the nematode EV samples were between 15 and 25 nt in length (Figure 5A). The *C. elegans* microRNA FASTA file was downloaded from miRBase version 21^(Kozomara and Griffiths-Jones, 2014).^ FASTA files for *C. elegans* transcriptome and genome sequences were retrieved from NCBI genome database (assembly WBcel235). The FASTA files were transformed with bowtie-index files. All the alignments to ncRNAs, RNAs and DNAs were performed under 0 to 2 mismatch allowance. This resulted in ~14M total reads of which ~11.5M were mapped to various sequence databases with 0, 1, or 2 mismatches. Among the mapped reads, 8.5M of these reads mapped to the *C. elegans* sequences with 0, 1, or 2 mismatches while 2.5 M were from *E. coli* sequences (Figure 5B). These *C. elegans* hits were then categorized as originating from the plus or minus strand. The most abundant RNA species by far was rRNA, comprising over 6M of the total reads (Figure 5C). We then filtered out these rRNA reads and determined the relative abundances of the remaining RNA species limiting our analysis to transcripts with no-mismatches. This resulted in 440,392 reads. The distribution of non-rRNA types and species within a type are summarized in Table 2.

**Figure 5:**
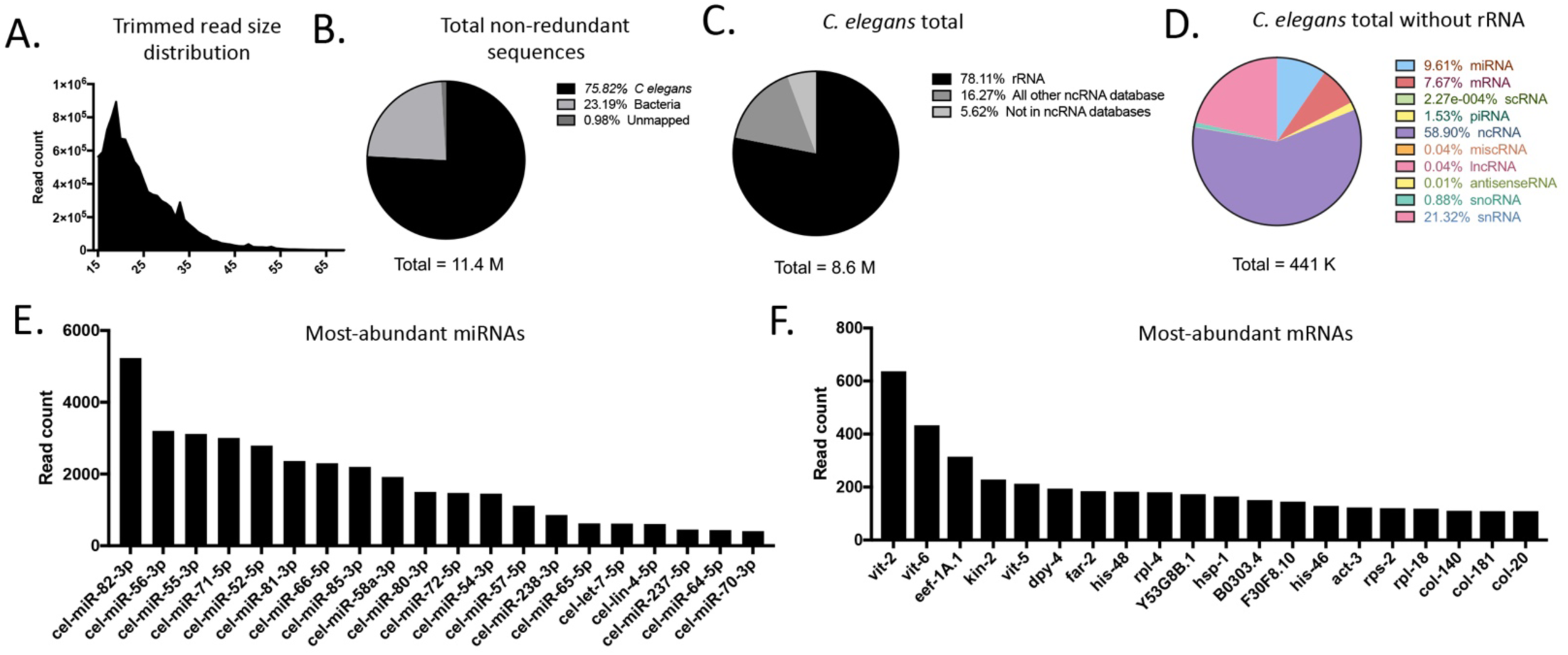
RNAseq analysis *of C. elegans* EVs reveals different classes of RNA: A) Size distribution of high-quality RNA reads The majority of the reads were between 15 to 15 nt in length B) Distribution of three major categories of total non-redundant reads with up to 2 nucleotide mismatches. C) Distribution of total non-redundant C. *elegans* reads with no mis-matches. D) Distribution of species of non-coding RNA mapped with no nucleotide mismatches E) Top 20 miRNAs identified with no nucleotide mismatches F) Top 20 mRNAs identified with no nucleotide mismatches.

We then filtered for sequences that had no mismatches and identified the most-abundant species within each class of RNA. The top 20 most abundant miRNA species included miR-80-82, 85 as well as miR-55-57, miR-70-72, as well as let-5 and lin-4 (Figure 5F). The top 20 mRNA transcripts included vitellogenin, collagens, ribosome and histone species, as well as translation elongation factor eef-A.1, the HSP70 encoding transcript *hsp-1*, and several uncharacterized genes (Figure 5G). The top reads for the other species are shown in supplemental and the filtered BAM file for the sequencing is publicly available (Supplemental Figure 7).

## DISCUSSION

Here we provide the first report on isolation, purification, and -omics characterization of environmentally-secreted EVs from *C. elegans*. To achieve this, we established new methods for isolating EVs and performing analyses typically used for studying mammalian EVs, including nanoparticle tracking analysis, flow cytometry, proteomics, and RNA sequencing.

Our proteomic results indicate that *C. elegans* EVs are specialized organelles with protein cargoes distinct from the spectrum of proteins found in whole worm protein lysate and possess many of the EV cargos most-identified in mammalian EV proteomic studies. The set of proteins detected from worm EVs was highly differentiated from the most abundant proteins in whole worm lysate with the most-abundant EV proteins showing very significantly higher normalized spectral abundance values. This suggests that the protein cargo loading of *C. elegans* EVs is an active and selective process, raising the possibility for using the genetic power of *C. elegans* to study the cellular mechanisms that regulate EV cargo loading.

We uncovered that *C. elegans* EV proteins are highly-enriched for membrane raft proteins, which have been shown to serve in membrane organization and are characteristic of mammalian EVs ^(Lingwood and Simons, 2010)^. The lipophilic dye (DI-8-ANEPPS) used in our FACS experiments binds to cholesterol, one of the main components of lipid rafts. Our proteomics results overlap considerably with previous analysis of the *C. elegans* lipid raft proteome extracted from whole worms ^(Rao et al., 2011)^. 25 of the 43 genes identified in their lipid raft proteomics study also were also present in our EV proteomic data sets (Supplemental Table 2). The total worm lipid raft proteins are associated with several different cellular compartments. The majority (7 of 10) of the total worm lipid raft proteins with known associations to “membrane” or “extracellular” were identified in our samples (Supplemental Table 2). Because lipid raft proteins are membrane markers by definition, and some of these likely are exposed to the outside of the EV this set of 25 proteins likely contains EV protein markers that can be adapted with extracellular-facing small affinity tags for immunoprecipitation of tissue-specific *C. elegans* EVs.

EVs are hypothesized to interact with the dense polysaccharides on the exterior surface of the cell membrane facilitating their uptake into recipient cells ^(Mulcahy et al., 2014),(Escrevente et al., 2011)^. Our gene ontology analysis identified that *C. elegans* EVs are enriched for proteins that interact with polysaccharides. The most enriched protein domain was the “lectin C-type domain”, known for polysaccharide binding and the next most-enriched was “transmembrane glycoprotein”. Our proteomics results also identified PAT-3 with strong orthology to human integrins that interact with the glycoprotein fibronectin matrix on the exterior surface of cells ^(Hagedorn et al., 2009)^. This suggests *C. elegans* may be a useful genetic model for studying extracellular receptor-ligand binding in the context of EV signaling. For instance, receptor ligands interactions could be studied in a cell-specific manner by fusing split-GFP ends to receptors and ligand proteins in different cell types.

In addition to abundant rRNA, mammalian EV RNA cargo include messenger RNA (mRNAs), non-coding RNA (ncRNAs) including miRNAs, long non-coding RNA (lncRNA), single-stranded DNA (ssDNA), double-stranded DNA (dsDNA), mitochondrial DNA, and oncogene amplifications (i.e., c-myc) ^(Janas et al., 2015),(Thakur et al., 2014),(Guescini et al., 2010),(Yáñez-Mó et al., 2015)^. It was not surprising that a majority of the mapped reads corresponded to ribosomal RNA because these species are known to be highly abundant in all cells and packaged into EVs ^(Jenjaroenpun et al., 2013),(Miranda et al., 2010)^. For this reason, RNA extraction procedures have been developed to reduce their numbers to allow for identification of other RNA types in RNAseq experiments. Our RNAseq analysis gave robust results with > 440,000 reads mapped to other RNA species, including ncRNA, mRNA, miRNA, piRNA, and snRNA among others with perfect match (Figure 5D). Non-coding RNA was the most abundant species of RNA identified comprising 58% of the non-rRNA reads. Almost a third of the ncRNA reads were for three uncharacterized species, M02F4.12, C44B7.15, and B0244.13. Although little is known about small ncRNA functions, the enrichment of small ncRNAs in EVs suggest they may play a role in cell-to-cell signaling. Small non-coding RNA and rRNA has been shown to also be enriched in human exosomes ^(Nolte-’t Hoen et al., 2012)^. MicroRNA was the second most-abundant family of RNA comprising ~10% of the non-rRNA reads. The reads mapped to 139 miRNA species which are all encoded from the plus strand. The uncharacterized microRNA miR-82 had the most reads by far (Figure 5F). Intriguingly, one of the most abundant microRNAs was let-7 which has been recently been identified in human cancer cells and breast milk EVs ^(Ohshima et al., 2010),(Kosaka et al., 2010)^. The piRNAs identified were notable because of their great variety, 6,776 reads mapped to >1,200 different transcripts. In contrast, we only identified 139 miRNA species from 42,444 miRNA reads without mismatches. One of the advantages of RNAseq analysis is that different small RNA families can be studied simultaneously in a comprehensive manner. Through the methods described here RNAseq analysis can now be applied to *C. elegans* EVs.

It remains an open question whether all cell types secrete EVs or if EV secretion is restricted to a subset of cells. Although it was shown by Wang *et. al*. that EVs from ciliated sensory neurons are secreted outside the body of the nematode, it seems likely that other tissues such as the intestine and excretory canal also produce EVs. Our mass spectrometry data suggest that there are EVs present in secretate that derive from many different kinds of tissues, including neuronal, intestine, gonad, and muscle. As all tissues can empty into the pseudocoelomic space which is filtered into the excretory canal, it is feasible that all cells may have the capacity for transmitting EVs outside of the animal. EV signaling and excretory functioning in *C. elegans* may be physiologically associated because they have both been shown to be influenced by the functions of similar genes, including *vha-5*, a vATPase, and *ral-1* a small GTPase ^(Liégeois et al., 2006),(Armenti et al., 2014)^. With the identification of numerous EV membrane-marking proteins in this study, it should be possible to generate tissue-specific EV-markers for identifying which tissues contribute to the total pool of secreted EVs.

## Limitations and opportunities for future development

The methods we developed result in abundant, relatively pure EVs sufficient for conducting multiple types of analysis. However, we note that in order to separate the worms from bacterial contaminants, the animals were incubated in buffer completely free of bacteria for a period of time prior to collection of EVs. We determined that the animals were healthy and active following the incubation in S basal, suggesting that no death or severely adverse effects occurred. However, the EVs we obtain may reflect a starvation response that could influence the cargos and composition of the purified EVs. Future effort will be placed toward purification of secreted EVs from *C. elegans* without washing and incubation in S basal; however, as bacteria are vastly more numerous than the nematodes, when EVs are purified from well fed worms using our current methods we find that bacterial peptides comprise ~ 90% of the total peptide hits in proteomic experiments. This reduces the ability to identify lower-abundance nematode EV peptides. By identifying robust EV membrane protein markers, we anticipate that it will be possible to develop one-step immunoaffinity purification methods similar what is currently being conducted with mammalian EV studies.

Although human anti-CD63 conjugated dye marked detergent-labile vesicles in our FACS analysis, we did not uncover the *C. elegans* orthologs of commonly identified tetraspanin EV markers in our proteomics experiments. These tetraspanin markers are frequently identified in human EV proteomics studies because human EVs are often immunoprecipitated with antibodies to these proteins, therefore the EV populations under study are enriched for these markers even if the total EVs from the original experimental sample was not enriched for tetraspanins. Our proteomics was sufficient for identifying the most abundant proteins in EVs, but we recognize that it was not a comprehensive analysis of all *C. elegans* EV protein cargoes. In these experiments we analyzed the total protein of EVs and therefore were biased towards soluble proteins cargos due to their greater abundance than transmembrane proteins. In the future it will be informative to process EVs samples with a protein extraction kit that enriches for membrane proteins ^(Qoronfleh et al., 2003)^. Greater proteomic coverage can also be obtained through increasing the biomass of the EV samples, extending the elution times, and cutting out each band in the Coomassie gel in for separate digestion and runs. With these experimental adjustments more comprehensive and quantitative protein measurements membrane marking proteins can be obtained so low-abundance markers specific to EV-subclasses could be identified.

The focus of our RNAseq analysis was to determine whether *C. elegans* EVs contained small RNA cargos, and if so, to develop a purification and sequencing pipeline that produces robust mapped reads. The abundant mapped reads (> 8 million with up to 2 mismatches, > 440,000 with no mismatches) and identification of the major classes of RNA contained in *C. elegans* EVs is sufficient to conclude that multi-omics analysis of *C. elegans* EVs is now a feasible approach. This opens the possibilities for exhaustive analytical characterization studies in the future. Through characterizing multiple biological replicates in which the RNA was physically size-selected prior to building the RNAseq library and including internal RNA standards it will be possible to obtain higher coverage of small RNA species and conduct sensitive comparative analysis between experimental samples.

## Conclusion

EVs are involved in virtually every aspect of human health and disease. The development of optimized isolation, characterization and quantification of EVs from *C. elegans* is a significant step forward towards leveraging this powerful invertebrate genetic model for better understanding the compositions and signaling properties of EVs under different physiological conditions. In this report we found that *C. elegans* EVs share many properties of mammalian EVs, including characteristic protein cargos and RNA species, membrane composition, and transmembrane marker proteins. These results suggest that *C. elegans* EV-signaling has functional evolutionary conservation with humans. The genetic strength of *C. elegans,* coupled with the ability of isolate EVs on a scale amenable to FACS sorting and multiple parallel downstream analysis, suggest that this simple nematode could be a powerful model for multi-omic cargo analysis of EVs.

## METHODS

### Strains

For this study we utilized wild type (N2) *C. elegans*. Worms were cultured and maintained using standard methods ^(Brenner, 1974)^.

### Generation and purification of EVs

Synchronous populations of animals ~500,000 animals were grown to young adulthood at 20C on HGM media with NA22 bacteria at a density of 20,000 per 10 cm plate to allow them to reach adulthood without starving. Upon reaching young adulthood the animals were washed off the plates with S. Basal buffer and put into a 50 mL conical polypropylene tube. The animals were allowed to settle by gravity for 5 minutes and the supernatant was removed. The animals were then combined and washed five more times with S. Basal. The animals were then suspended in 30% ice-cold sucrose cushion and centrifuged at 1000 X G for 3 minutes. The floating animals were transferred into a fresh 50 mL conical tube, and washed 3X with 50 ml S. Basal. After the third wash the animals were suspended in S. Basal + 2.5 μg/mL cholesterol at a density of 1 animal per μL in 50 ml conical tubes filled up to 45 mL. These tubes were then incubated at 20C in an end-over-end rotator for 24 hours. At the end of the incubation period the animals were pelleted at 2500 X G for 10 minutes and the supernatant transferred into a fresh 50 mL conical tube. Pellet any remaining worms at 2500 X G for 10 minutes and then pass supernatant through a 0.2 μm filter. Any remaining larger lipid particles were then pelleted at 18,000 X G for 30’ at 4C. The supernatant was decanted and then concentrated over a 10 kD regenerated nitrocellulose filter (Amicon Ultra-15) by centrifuging at 2500 X G until the volume is reduced to 1500 μL. Protease inhibitor cocktail (HALT + EDTA) was added to the concentrated sample. The concentrated fraction was then passed over a 10 mL Sepharose CL-2B size exclusion column with a mobile phase of S Basal + protease inhibitor cocktail at 4C. 15 mL of elute was collected in 1 mL fractions. After characterizing the EVs through TEM we determined that if we combine the elution volumes 2-6ml we capture most of the EVs but almost no soluble proteins. Thereafter it became standard procedure to consolidate the 2-6 mL, 7-11 mL, and 12-16 mL elution fractions and then concentrate over a 10 kD regenerated nitrocellulose filter to a final volume of 100 μL. Samples were then either analyzed immediately or stored at −80C until ready for downstream applications. Total protein of *C. elegans* secretate fractionation was determined on a QuBit 4 fluorometer with a QuBit protein assay kit (Thermo Fisher Rockford IL, USA Cat #s Q33226, Q33211).

### Nanoparticle tracking analysis

The SEC fractions were analyzed via nanoparticle tracking analysis (NTA) by standard methods using a NanoSight ns3000 (Malvern Panalytical, Worcestershire UK). Briefly, each sample were diluted with 0.2 μm filtered MilliQ water until 20 −100 particles were identified in the field of view (generally 1:100). In NTA the paths of particles act as point scatterers, undergoing Brownian motion in a 0.25-ml chamber through which a 532-nm laser beam is passed, is determined from a video recording with the mean squared displacement determined for each possible particle. The diffusion coefficient and sphere-equivalent hydrodynamic radius are then determined using the Stokes–Einstein equation, resulting in a distribution of particle size. NTA 3.1 software was used for particle analysis. The consolidated size exclusion fractions (2-6 mL, 7-11 mL, 12-16 mL) were diluted 1:100 in 0.22 μm-filtered S Basal was run as the buffer control. For a negative control the S. Basal buffer itself was run by itself. The settings for the acquisition were as follows.

Camera level 14, slider shutter 1259, slider gain 366, 30 ms exposure, 25 frames per second, particle detection threshold 5, max jump distance 11.6 pixels. Three independent 60 second acquisitions were obtained for each sample. Five biological replicates were quantified. This resulted in an average of 55 verified particle tracks in each frame of the video with a standard deviation of 15 particles and greater than 700 particle tracks per 60 second movie.

### Transmission electron microscopy

We glow discharged formvar-carbon coated copper mesh grids (Polysciences, Warrington, PA USA; Cat # 24915-25) with a hydrogen/oxygen mix for 30 seconds on a Gatan Solarus 950 Plasma Cleaner (Gatan, Pleasanton, CA USA). To analyze our size exclusion column elution we spotted 2 μL of the fractions onto glow discharged formvar-carbon coated copper mesh grids (Polysciences, Warrington, PA USA; Cat # 24915-25) incubated for 2 minutes, wicked the sample off with Whatman 2 filter paper (Millipore Sigma, Burlington MA, USA; Cat # WHA1002055), stained with 2% phosphotungstic acid adjusted to pH 7.0 (PTA) (Ted Pella Redding, CA USA; Cat # 19402) and washed the grids three times with 2 μL filtered MilliQ water. We imaged our samples at 19,000 X with a Philips CM100 TEM at 80kV.

### Flow cytometry

Three biological replicates were analyzed using an Apogee A50 flow cytometer (Apogee Flow Systems). Flow cytometer sheath solutions were 0.1 μm filtered before use. Polystyrene fluorescence beads (0.11 and 0.50 Km) and size calibrated non-fluorescent silica beads (0.18, 0.24, 0.30, 0.59, 0.88, and 1.30 μm) were resolved with forward (Small Angle Light Scattering) and side (Large Angle Light Scatter) light scatter (Fig.1A); 0.18 um was the lower limit for size resolution of beads (Supplemental Figure 2). Because vesicles have a different refractive index than the beads they run differently on the FACS. Therefore, we did not use the calibration beads as a means to quantify absolute size of the particles but simply as a way to check the consistency of the Apogee A50 performance before analyzing our experimental samples. We analyzed the volume of each sample by running each at 1.5 μL/min for 180 seconds. Analysis was conducted with FloJo software (FloJo, Ashland, Oregon USA). To determine dye-labeled particles we gated from SALS level 100 to 10000 (~100 nm to 300 nm sized particles) on the fluorescence intensity such that the iso-type sample with no dye give a reading of 2.5% of the total events. We then assessed the gate on the dye-only at the same concentration as we used for our experimental samples. To determine if the particles were detergent sensitive we added Triton-X 100 to the sample for a final concentration of 0.05%. We then sonicated the sample gently at an intensity level three with a 30% duty-cycle for 10 cycles before analyzing on the Apogee A50.

### Proteomics

We prepared three biological replicates of EVs from young adult N2 animals as above. Total protein in the EV fractions were quantified by QuBit analysis and samples were normalized for protein concentration. These were then extracted in RIPA buffer with vortexing for 30 minutes before adding NuPage 4X sample buffer and heating to 70 C for 10 minutes. Samples were then spun at 18kG for 15 minutes to pellet any precipitates. 50 μg of protein was loaded onto a large volume capacity SDS-PAGE gel and run at 80 mV for 10 min and then 100 mV until the dye front was ~ 1 cm below the start of the resolving gel. The gels were then washed twice with 250 dH_2_O for 60 minutes to remove any detergent. The samples were then cut out of the gel with a new razor blade and kept in a low-protein binding eppi-tube (ThermoFisher, San Jose, CA USA, Cat # 90410).

### In-Gel Digestion

Gel bands were washed in 100 mM ammonium bicarbonate, reduced with DTT, and alkylated with IAA. Gel bands were then shrunk with acetonitrile and speed-vacuumed to dry the gel bands. The gel bands were then digested with 1 μg of trypsin overnight at 37°C with shaking. The next day proteins were extracted from gel bands with 60% acetonitrile, 0.1% trifluoroacetic acid, and then reconstituted in 0.1% formic acid.

### Liquid Chromatography and Mass Spectrometry

Fused silica microcapillary columns of 75μm inner diameter (Polymicro Technologies, Phoenix, AZ) were packed in-house by pressure loading 30 cm of Repro sil-pur C18 material (Dr. Maisch, Gmbh, Germany). Kasil (PQ Corporation, Malvern, PA) frit microcapillary column traps of 150μm inner diameter with a 2 mm Kasil frit were packed with 4 cm of Repro sil-pur C18 material. A retention time calibration mixture (Pierce, Rockford, IL) was used to assess quality of the column before and during analysis. Three of these quality control runs are analyzed prior to any sample analysis and then after every six sample runs another quality control run is analyzed. Samples were loaded onto the trap and column by the NanoACQUITY UPLC (Waters Corporation, Milford, MA) system. Buffer solutions used were 0.1% formic acid in water (Buffer A) and 0.1% formic acid in acetonitrile (Buffer B). The 60-minute gradient of the quality control consisted of 30 minutes of 98% buffer A and 2% buffer B, 5 minutes of 65% buffer A and 35% buffer B, 6 minutes of 40% buffer A and 60% buffer B, 5 minutes of 95% buffer A and 5% buffer B and 18 minutes of 98% buffer A and 2% buffer B at a flow rate of 0.3 μL /min. The 180-minute gradient for the sample digest consisted of 130 minutes of 98% buffer A and 2% buffer B, 10 minutes of 60% buffer A and 40% buffer B, 1 minute of 40% buffer A and 60% buffer B, 6 minutes of 5% buffer A and 95% buffer B and 33 minutes of 98% buffer A and 2% buffer B at a flow rate of 0.3 μL /min. Peptides are eluted from the column and electrosprayed directly into a Velos Pro mass spectrometer (ThermoFisher, San Jose, CA) with the application of a distal 3 kV spray voltage. For the quality control analysis, a full-scan mass spectrum (400-1600 m/z) is measured followed by a 17 SRM spectra all at 35% normalized collision energy and a 2 m/z isolation window. For the sample digests, a full-scan mass spectrum (400-1600 m/z) followed by 17 data-dependent MS/MS spectra on the top 16 most intense precursor ions at 35% normalized collision energy with a 2 m/z isolation window. Application of the mass spectrometer and UPLC solvent gradients were controlled by the ThermoFisher XCalibur data system.

### Data Analysis

The quality control data was analyzed using Skyline ^(MacLean et al., 2010)^. The DDA MS/MS data was searched using COMET with dynamic modification searches of 15.994915 methionine and a static modification of 57.021464 Cysteine against a FASTA database containing all the protein sequences from the WS250 freeze of C. elegans from WormBase plus contaminant proteins ^(Eng et al., 2013)^. Peptide spectrum match false discovery rates were determined using Percolator ^(Käll et al., 2007)^ at a threshold of 0.01 and peptides were assembled into protein identifications using an in-house implementation of IDPicker ^(Zhang et al., 2007)^. The set was then filtered for proteins that were identified through two or more unique peptide hits, all protein isoforms condensed to the parent gene and the genes common to all three biological replicates were identified. This resulted in 161 high-confidence EV genes. This gene set was then analyzed for protein interaction and GO analysis with String.db.org ^(Szklarczyk et al., 2015)^. To determine how many of the top human EV proteins were also represented in our samples we downloaded the list of top 100 proteins most identified in ~500 proteomic studies that are uploaded to Exocarta ^(Keerthikumar et al., 2016)^. We found that 48 of the 100 proteins had reciprocal best hits in *C. elegans*. These 48 worm orthologs were then searched in our proteomics data resulting in 12 matches. A Fisher’s exact test was then applied to determine the significance of the overlap.

## RNA sequencing

### Sample preparation

The pipeline for RNA sample preparation and analysis is mapped out in Figure 5A. To quantify EV-associated RNA we processed our purified samples with Total exosome protein and RNA isolation kit (Invitrogen, Carlsbad CA, USA, Cat # 4478545). We then characterized the abundance and size of the elutes on an Agilent 2200 TapeStation (Agilent, Santa Clara CA, USA; Cat # G2991AA) using high-sensitivity screen tape (Agilent, Santa Clara USA Cat # 5067-5579) along with a small RNA calibration ladder (Agilent, Santa Clara CA, USA; Cat # 5067-1550). The vesicle samples displayed substantial small RNAs as well as 16s and 28s rRNA species at the expected sizes. To determine what RNA species are packaged into EVs we conducted RNAseq. We used a Qiagen small RNA Sample Preparation kit to prepare a cDNA library from our EV-associated RNA (Qiagen Germantown MD, USA Cat # 331502) while an Illumina MiSeq v2 kit (300 cycles) was used to prepare the sequencing library with 5’ adapter sequences (San Diego USA; Cat# MS-102-2002). The sample was then submitted to single-end sequencing on an Illumina MiSeq desktop sequencer.

### RNAseq data analysis

Adapter sequences were trimmed with Cutadapt and simple repeat sequences trimmed with PrinSeq. This resulted in RNA with a peak sequence length of 19 nucleotides (nt) dropping off to 0 reads around 55 nt. Adapter trimming and sequence alignment was conducted in the sRNAnalyzer ^(Wu et al., 2017)^. Sequences were searched against, miRBase ver 21 ^(Kozomara and Griffiths-Jones, 2014; Zerbino et al., 2018)^. FASTA files for *C. elegans* transcriptome and genome sequences were retrieved from NCBI genome database (assembly WBcel235) ^(Kozomara and Griffiths-Jones, 2014; Zerbino et al., 2018)^. The FASTA files were transformed with bowtie-index files. All the alignments to microRNAs, RNAs and DNAs were performed under 0 to 2 mismatch allowance. From a total of 14M reads, 11.5M were processed and 8M mapped to the *C. elegans* genome.

## Acknowledgements

We gratefully acknowledge Nick Terzopoulos for worm plates and reagents, the *Caenorhabditis* Genetics Center (CGC) for the N2 nematode line, Lucia Vojtech PhD for assistance with the nanoparticle tracking analysis, Jessica Young PhD and Marie Claire MD PhD for hiPSC-derived neuronal conditioned cell media, Safyie Celik for help with statistical analysis, Wai Pang for assistance with TEM imaging, and Albert Tai and Matthew Fierman of the Tufts University Bioseq facility for RNA sequencing. Sequencing was sponsored by the the BioSeq Program funded by the National Institutes of Health Science Education Partnership Award and the Cummings Foundation. This work was supported by NIH grant P30AG013280 to MK and NIH grant AG054098 to JCR.

## Author contributions

JCR conceived the study, carried out the experiments in the study unless otherwise indicated, and wrote the manuscript. GEM, JER and MJM conducted the mass-spectrometry experiments and analyzed the raw proteomics data to determine the statistically significant protein hits. NP and CDK advised on the FACS experiments and analysis, TK, KW, AN, and JCR analyzed the raw RNAseq data. MK supervised the study and wrote the manuscript.

---

**Figure S1:**
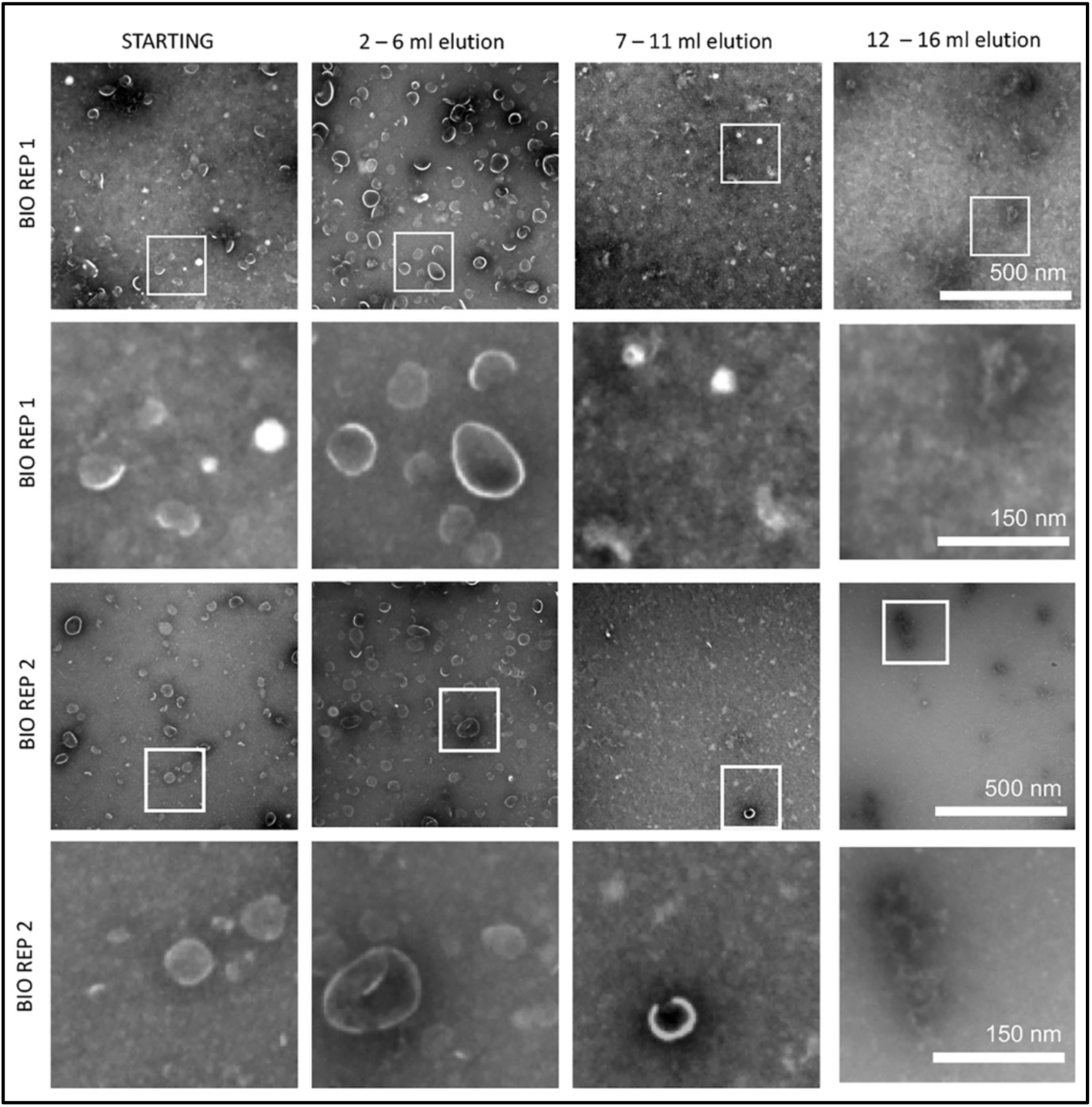
Transmission electron micrographs of size exclusion fractionated C. elegans secretate. Representative images of size exclusion elution fractions were negatively-stained and imaged at 19,000 X magnification. The 2-6 mL elution fractions contained the most particles with the EV cup-shape characteristic that EVs take on when dried and negatively stained. The white boxes indicate the area the area that is shown in the higher magnification images on the rows below. Scale bars shown on right.

**Figure S2:**
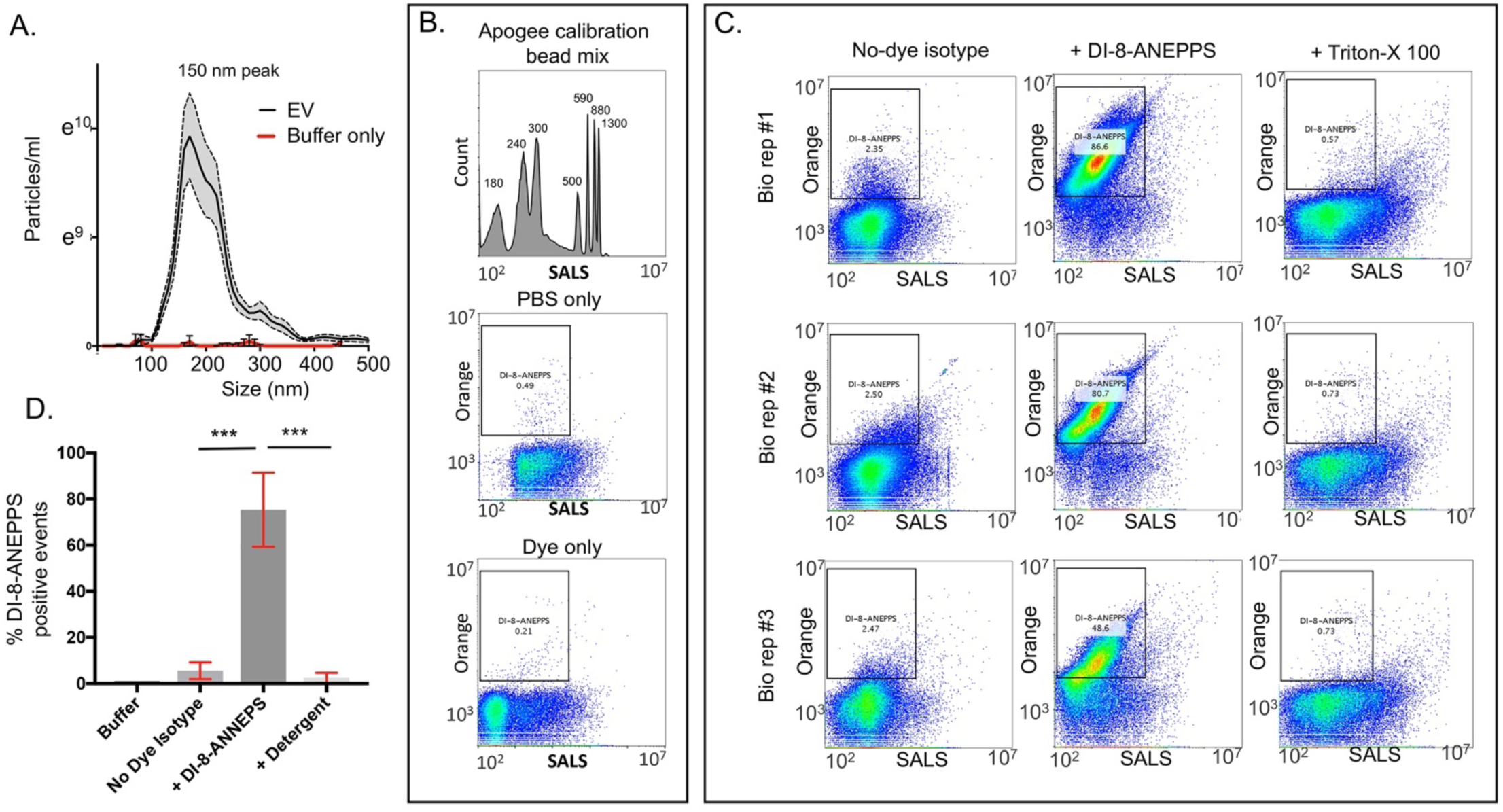
Verifying efficacy of EV purification scheme and FACS analysis with hiPSC-derived neuron EVs. Rationale: FACS analysis on a human cell culture EVs has been well established. Therefore, these experiments served as a positive control for our ability to effectively purify EVs and validate them as vesicles and not dense lipoprotein aggregates through labeling with DI-8-ANEPPS and then treating with detergent A) Nanoparticle tracking analysis of EVs purified from three biological replicates of hiPSC-derived neurons. B) Controls for flow cytometry analysis. On top is the histogram of the SALS profile of the Apogee Bead calibration mix. Peak sizes are indicated. Calibration bead mixes were run at the beginning of each day’s experiments to ensure the machine is separating sizes consistently. The buffer only and dye only isotype samples are shown below. The area used to gate for positive fluorescence events is shown. C) Three biological replicates of hiPSC-derived neuron EVs treated with DI-8-ANEPPS and detergent (0.05% Triton-X 100). The left column shows the no-dye isotype sample. The middle column shows the dye treated samples. The right column shows the dye treated samples subsequently treated with detergent. D) Summarized bar graph of the three biological replicates. Error bars in A and D are S.E.M. *** indicates a p-value of <0.001 as determined by a paired two-tailed T-test.

**Figure S3:**
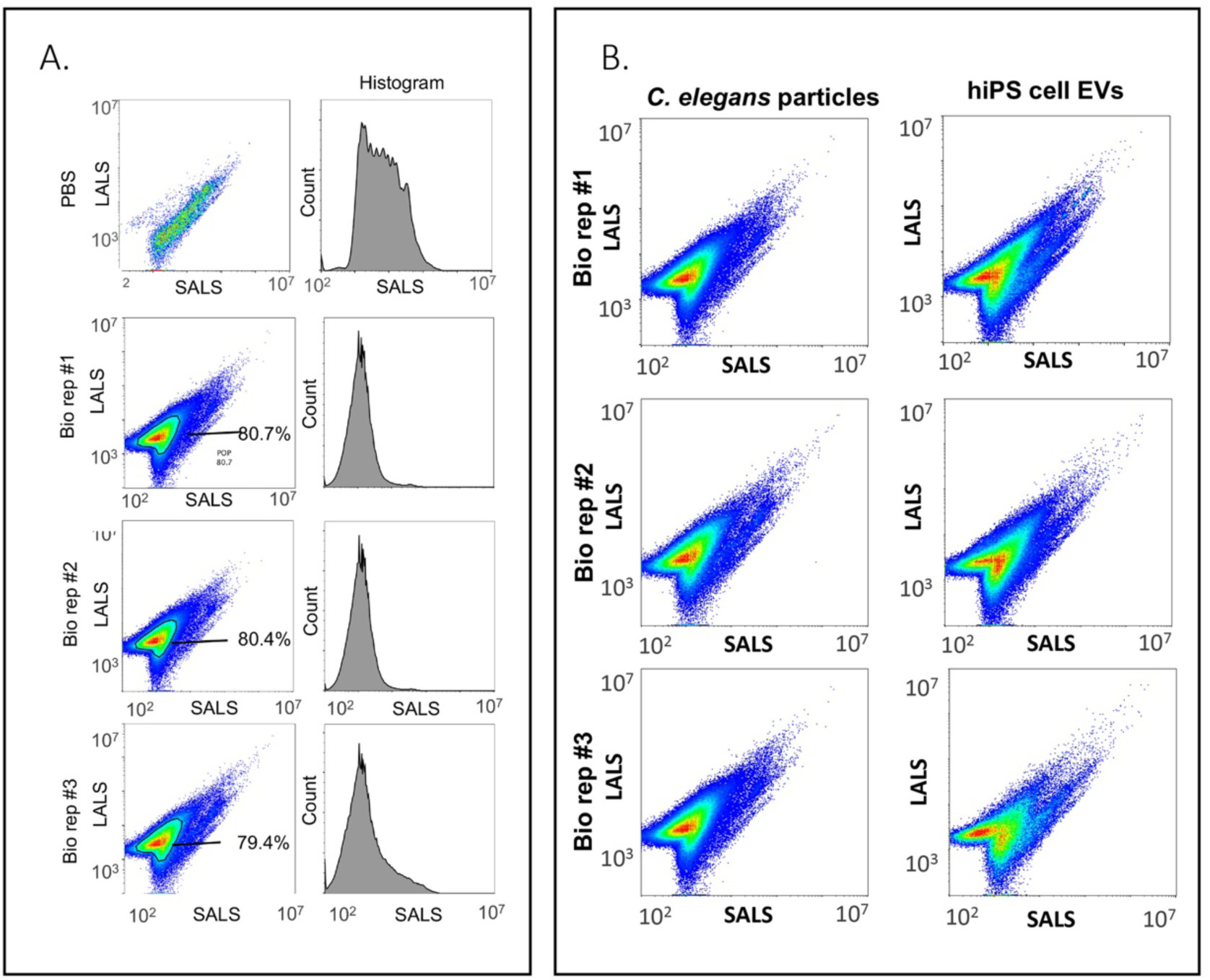
*C. elegans* EV preparations are monodisperse by small and large angle light scattering analysis. A) Histogram and SALS LALS plots of three biological replicates of purified *C. elegans* EVs. The distributions were similar. B) Comparison of SALS LALS scattering of *C. elegans* secretate EVs and hiPS-derived cell culture EVs.

**Figure S4:**
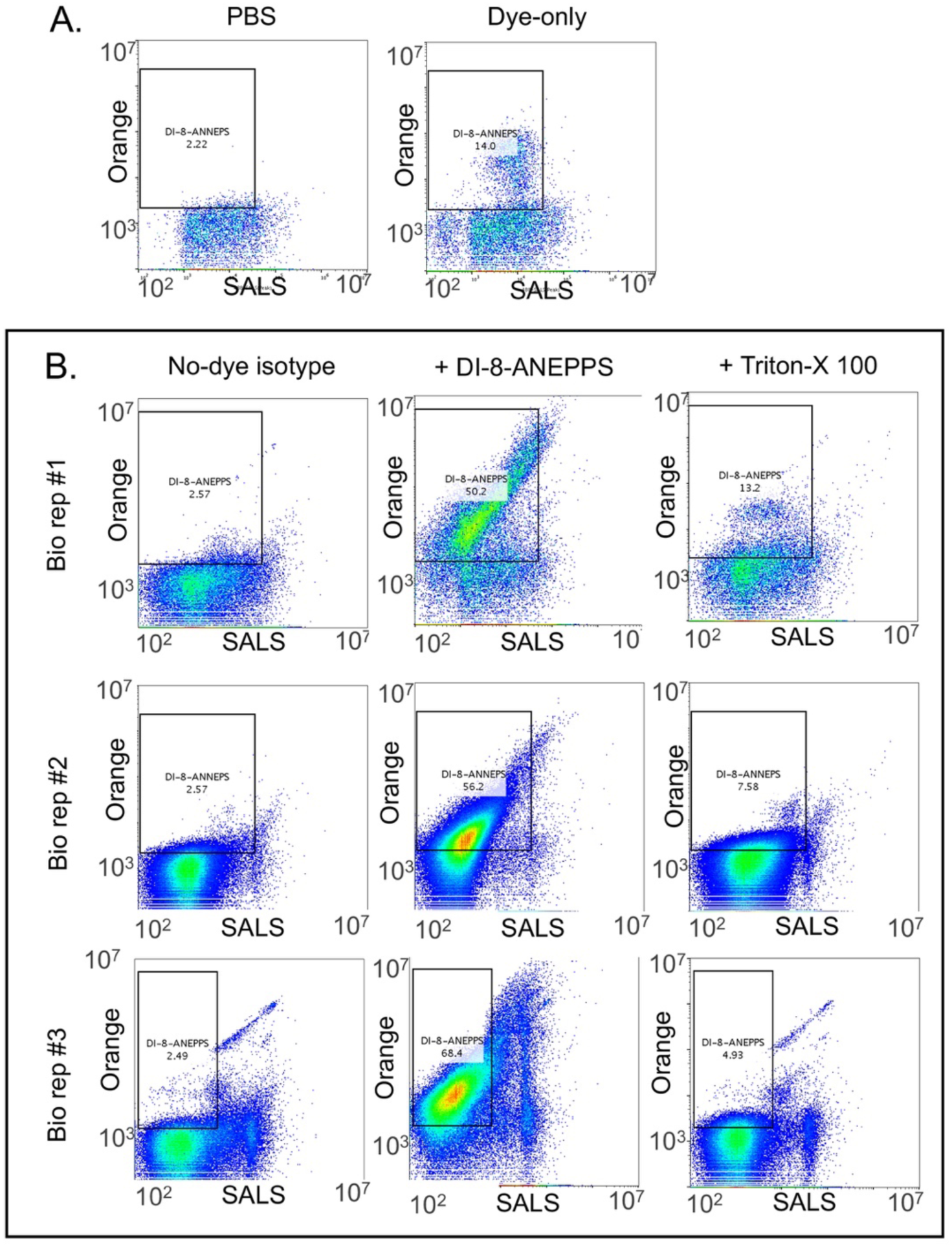
Particles purified from *C. elegans* secretate are extracellular vesicles and not solid lipoprotein particles. A) Buffer only and dye-only controls run under the same conditions (time, flow rate) as the biological samples. The box used to gate positive events is shown. Although the dye-only shows a high percentage of event the overall number is much lower than in the biological samples. (See Excel sheet in Supplemental) B) Three biological replicates of *C. elegans* EVs treated with DI-8-ANEPPS and detergent (0.05% Triton-X 100). The left column shows the no-dye isotype sample. The middle column shows the dye treated samples. The right column shows the dye treated samples subsequently treated with detergent. The dye treated samples show many positive events and these are eliminated by detergent treatment.

**Figure S5:**
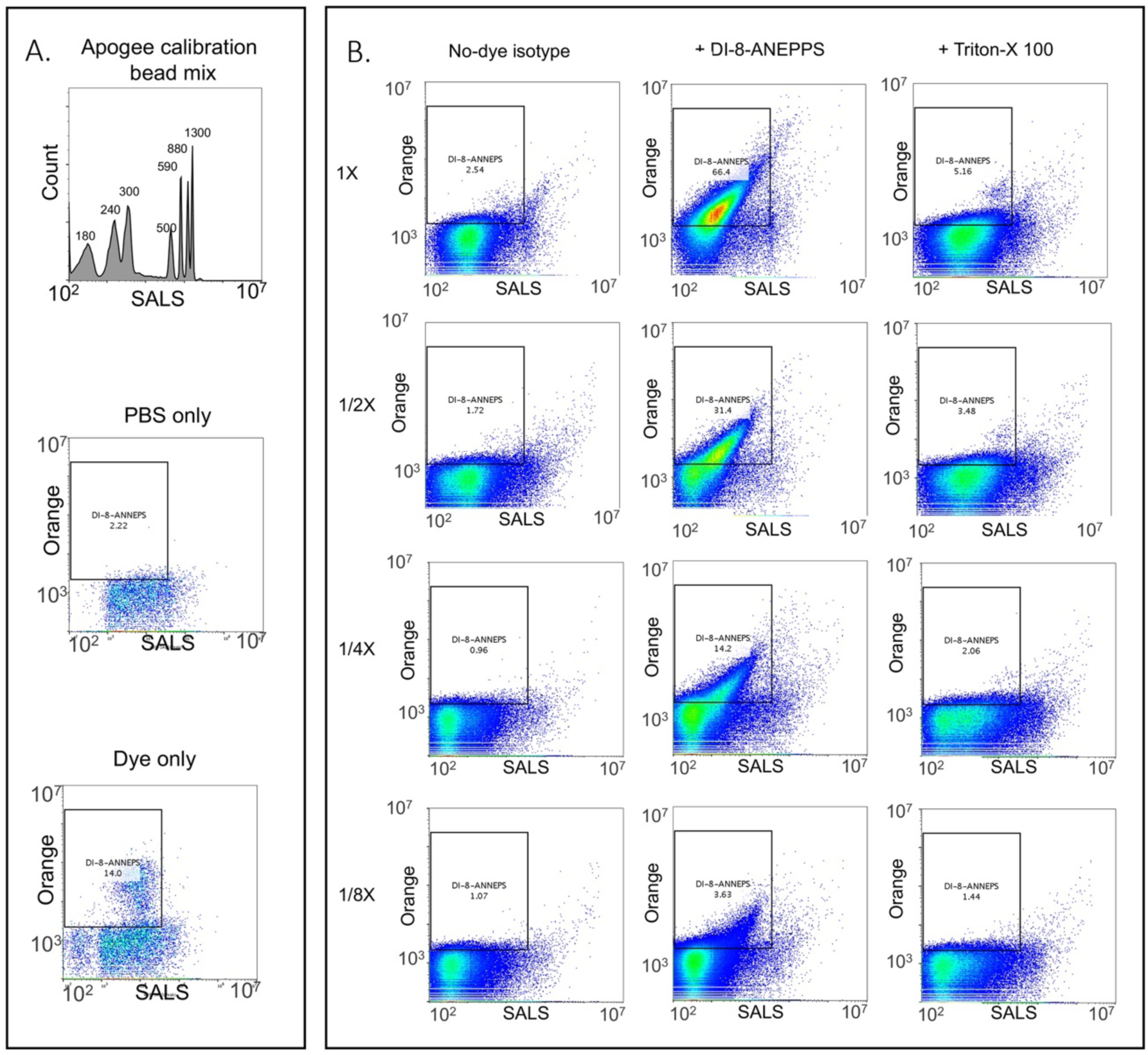
DI-8-ANEPPS-positive events are single vesicles. A) Controls for flow cytometry analysis. On top is the histogram of the SALS profile of the Apogee Bead calibration mix. Peak sizes are indicated. Calibration bead mixes were run at the beginning of each day’s experiments to ensure the machine is separating sizes consistently. The buffer only and dye only isotype samples are shown below. The area used to gate for positive fluorescence events is shown. B) Two-fold dilution series of DI-8-ANEPPS labeled EVs. The no-dye iso-type dilution series is on the left column, The DI-8-ANEPPS labeled sample is in the middle column and the detergent treated (0.05% Triton-X 100) is the left column. Each row is two-fold more diluted than the one above it. Note the percentage of gated events in the middle column decreases in proportion to the dilution.

**Figure S6:**
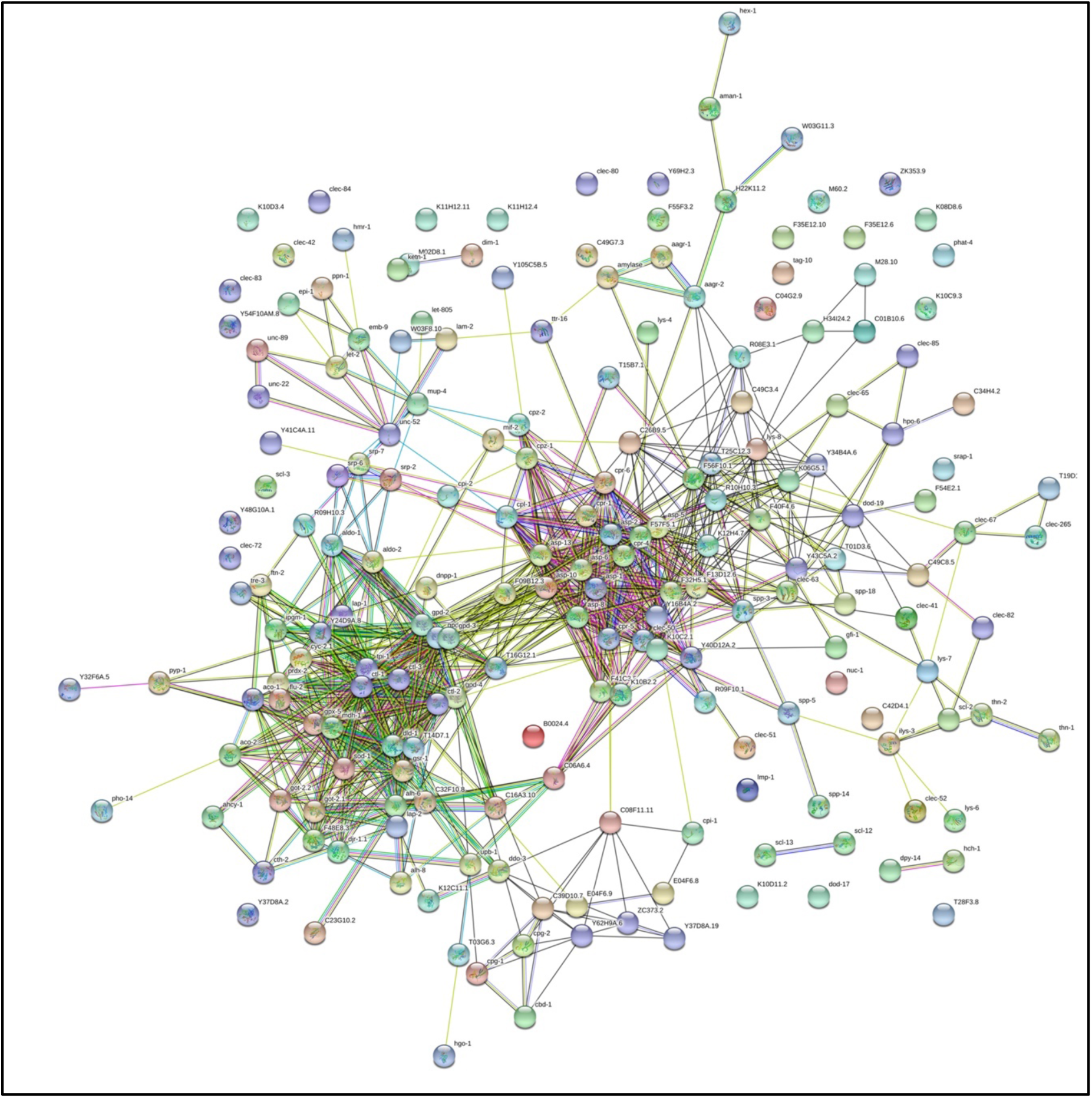
Protein interaction network of 161 proteins identified in all biological replicates. We analyzed all of the proteins that had one or more unique peptide hits in all our biological replicates with the protein interaction network analysis program String (Sting-db.org). The protein set showed two strong interaction clusters consisting of either protease genes or metabolic genes. There is also a small group at the bottom that is enriched for membrane raft proteins. 29 of the 161 proteins have no known interacting partners.

**Figure S7:**
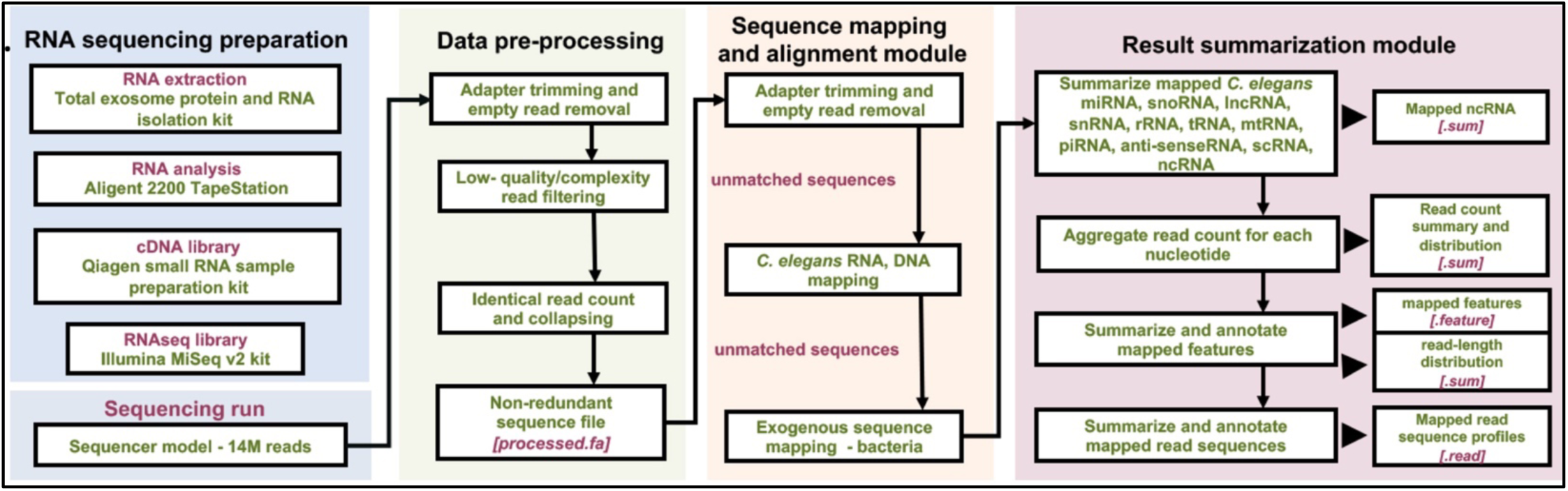
Analysis pipeline for identifying ncRNA from *C. elegans* EVs. We used the sRNAnalyzer pipeline for trimming adapter ends, filtering low-quality reads and collapsing identical reads. The reads were then iteratively mapped to 5’ or 3’ ends of miRNA, mRNA, and other ncRNA sequences (miRBase ver 21 and WBcel235. Mapped reads were calculated with 0, 1, and 2 mismatches allowed. The remaining unmatched sequences were then mapped to the genome.

**Figure S8:**
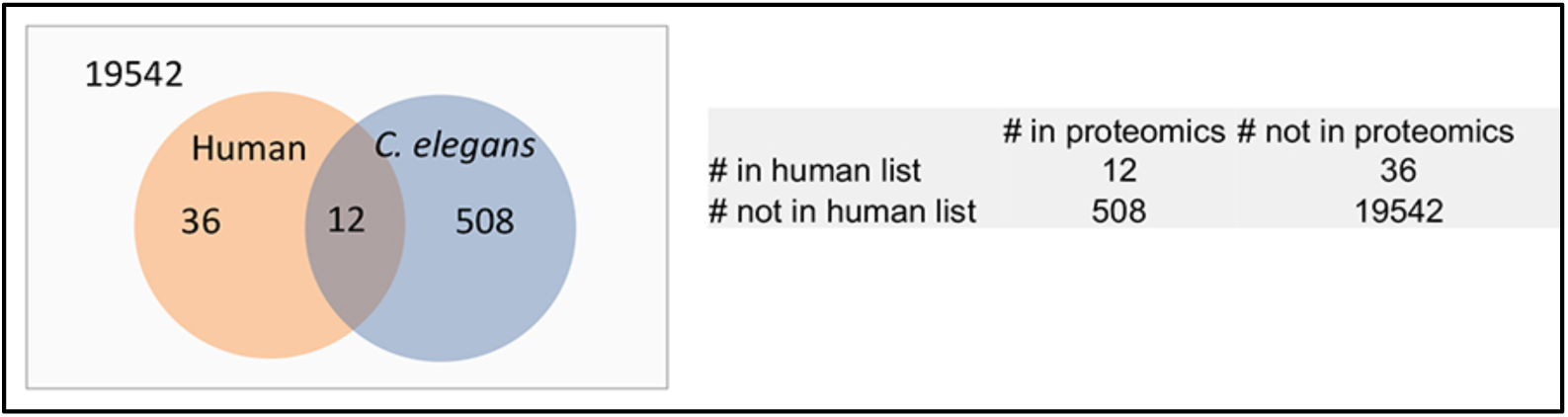
2 × 2 matrix for Fisher’s exact test to determine significance of overlap between C. elegans EV proteins and the most identified human EV proteins on Exocarta.org. 48 human top 100 most identified EV proteins had reciprocal best hits in *C. elegans*. Of these 12 of the proteins were found in *C. elegans* EV proteomics. A total of 520 proteins were identified with the proteomics. Calculations assume 20,000 protein coding genes in *C. elegans.* The P-value is 2.36^−9^ indicating that the *C. elegans* EV proteins overlapped with the human EV proteins much more than chance (Random chance = ~2 matches).

**Table S1:**
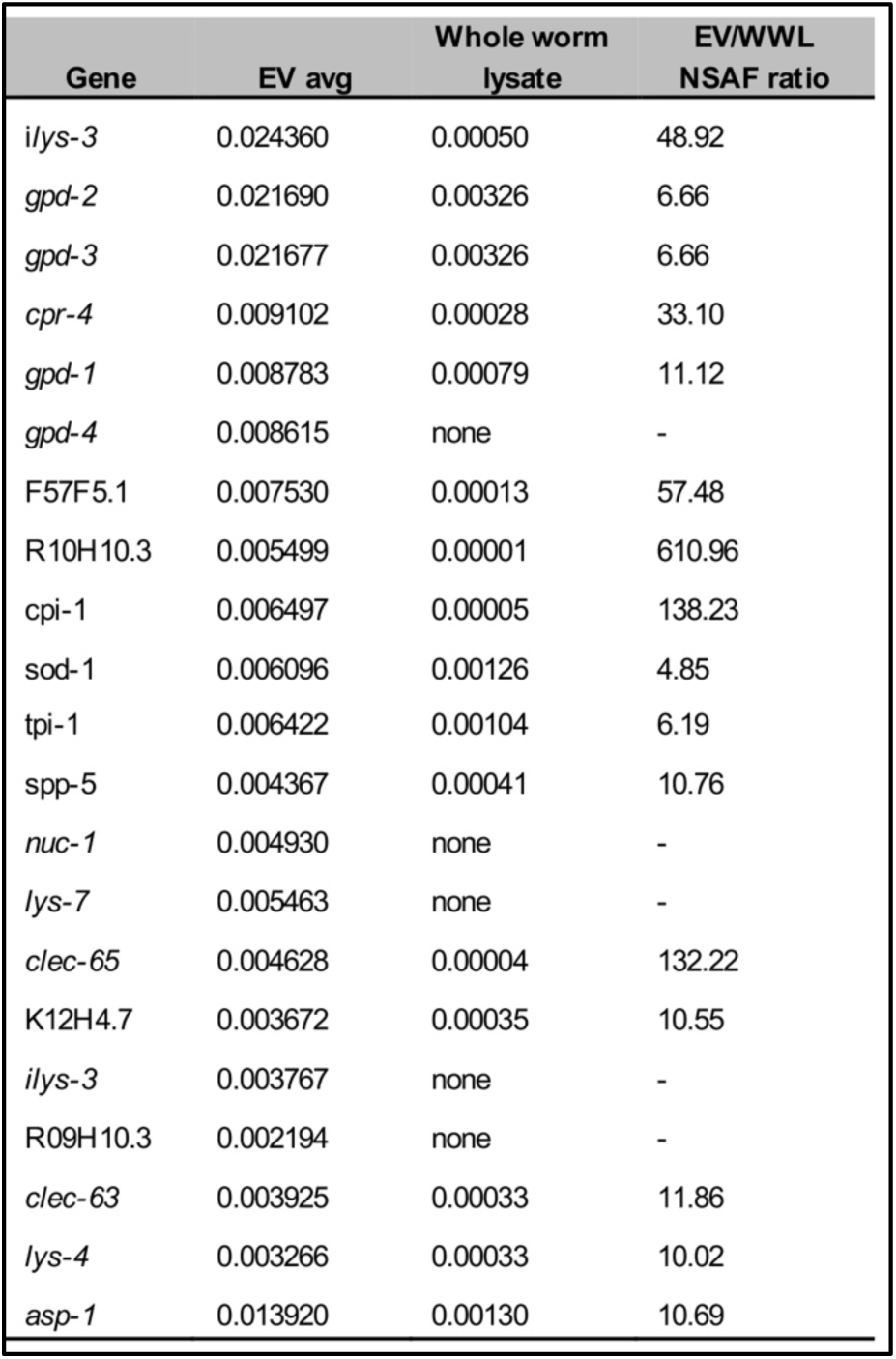
Comparison of normalized spectral abundance factor values top 20 EV proteins with their values from whole worm lysate. The top 20 most-abundant proteins observed in all three biological replicates were then compared with their values from whole worm lysate of day 1 adult animals obtained from a previous experiment ^73^. 25% of the proteins were not identified in whole worm lysate. All remaining 15 proteins had many-fold lower NSAF values in whole worm lysate. This indicates that the EV fractions contained a specifically enriched set of proteins and not just the most abundant proteins in the worm.

**Table S2:** EV samples are enriched for membrane raft proteins. A) 24 of the 161 (15%) of the EV proteins were associated with membrane rafts. This is over half of the 44 total *C. elegans* membrane raft proteins identified from whole worm lysate proteomics ^54^.

| Membrane raft proteins identified in EV samples | Cell compartment |
| --- | --- |
| K10C2.1 | Membrane |
| T19D12.4 | Membrane |
| F56F10.1 | Membrane |
| Y16B4A.2 | Membrane |
| *ctsa-2* | Membrane |
| *lec-4* | Extracellular |
| *lec-1* | Extracellular |
| *pho-1* | Brush boarder |
| *nep-17* | Lysosome |
| C26B9.5 | Lysosome |
| *gfi-1* | n/a |
| *hpo-6* | n/a |
| *lec-5* | n/a |
| *dod-19* | n/a |
| *tre-3* | n/a |
| K08D8.6 | n/a |
| *pcp-3* | n/a |
| F32A5.3 | n/a |
| F35E12.10 | n/a |
| *dct-17* | n/a |
| F54E2.1 | n/a |
| C29F3.7 | n/a |
| *tag-10* | n/a |

**Table S3:**
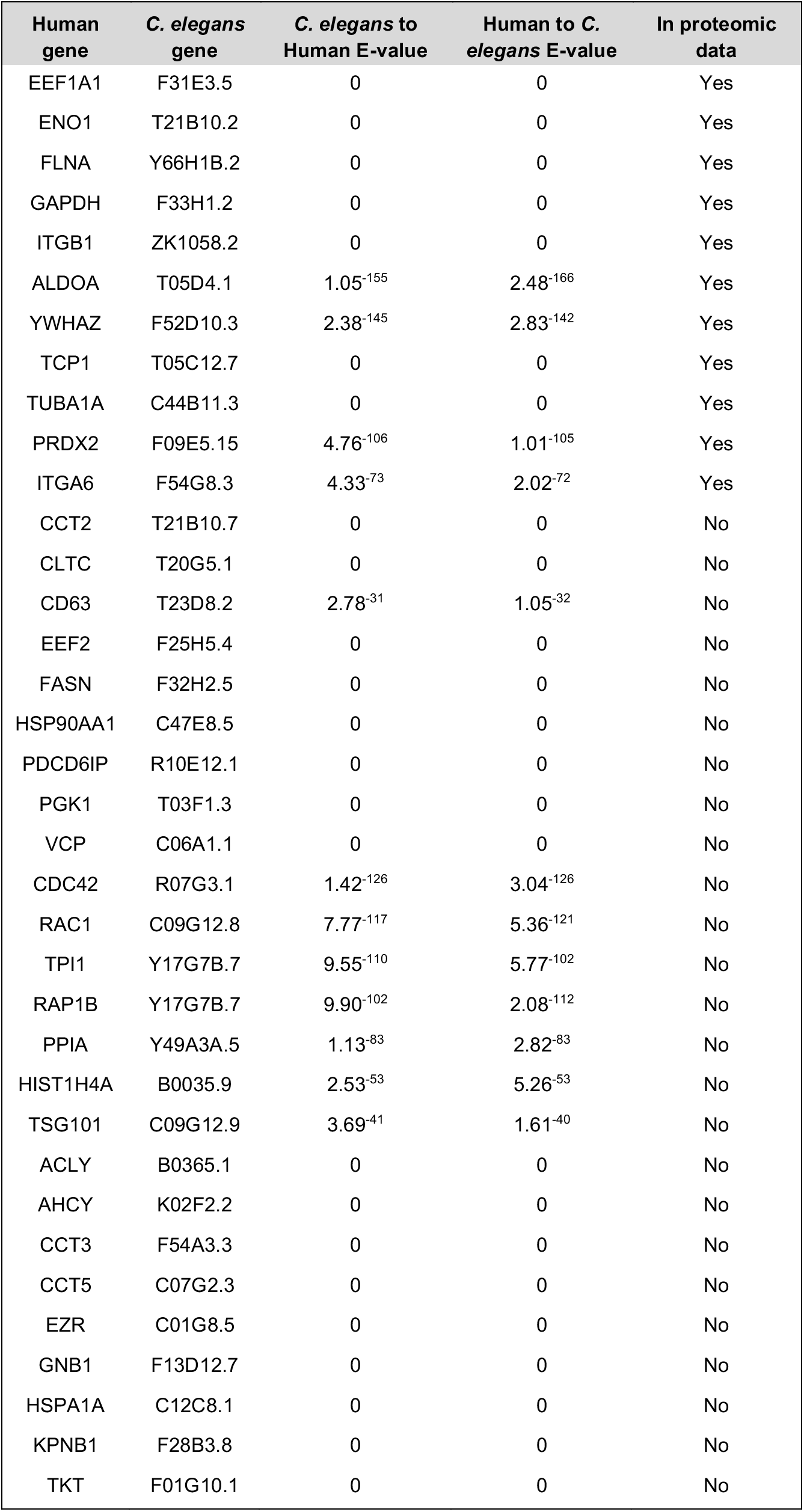
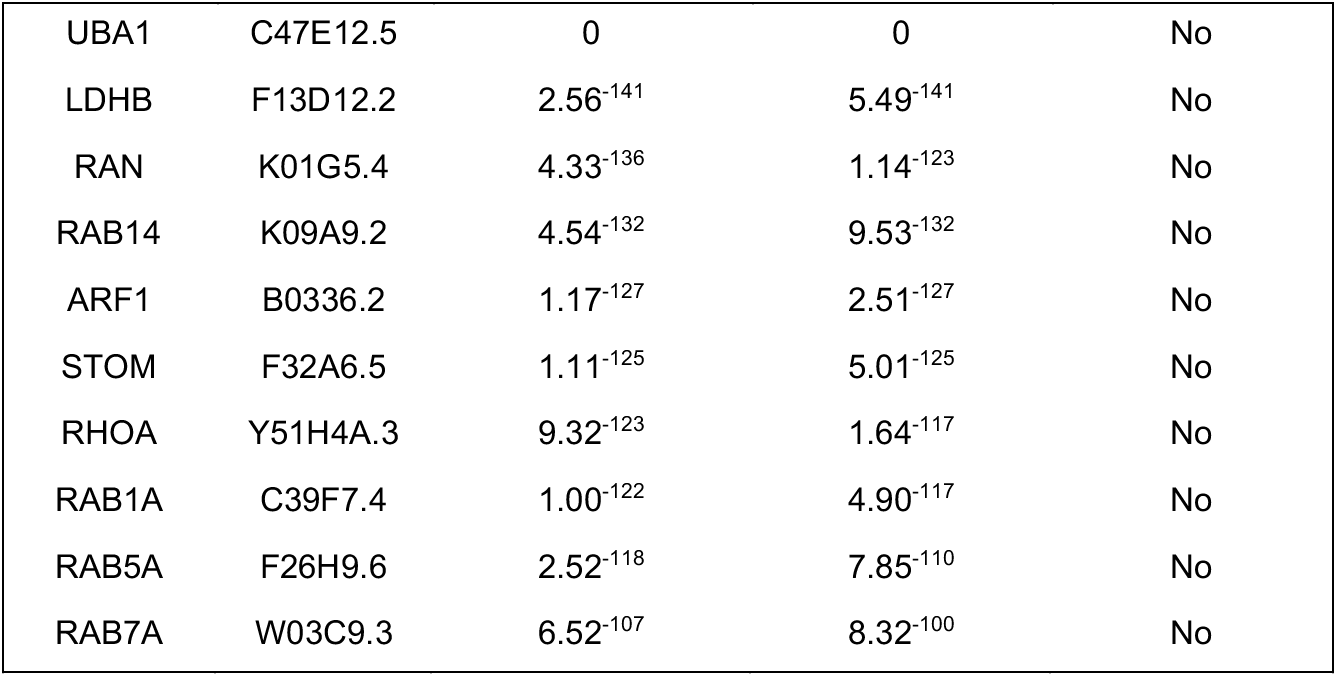
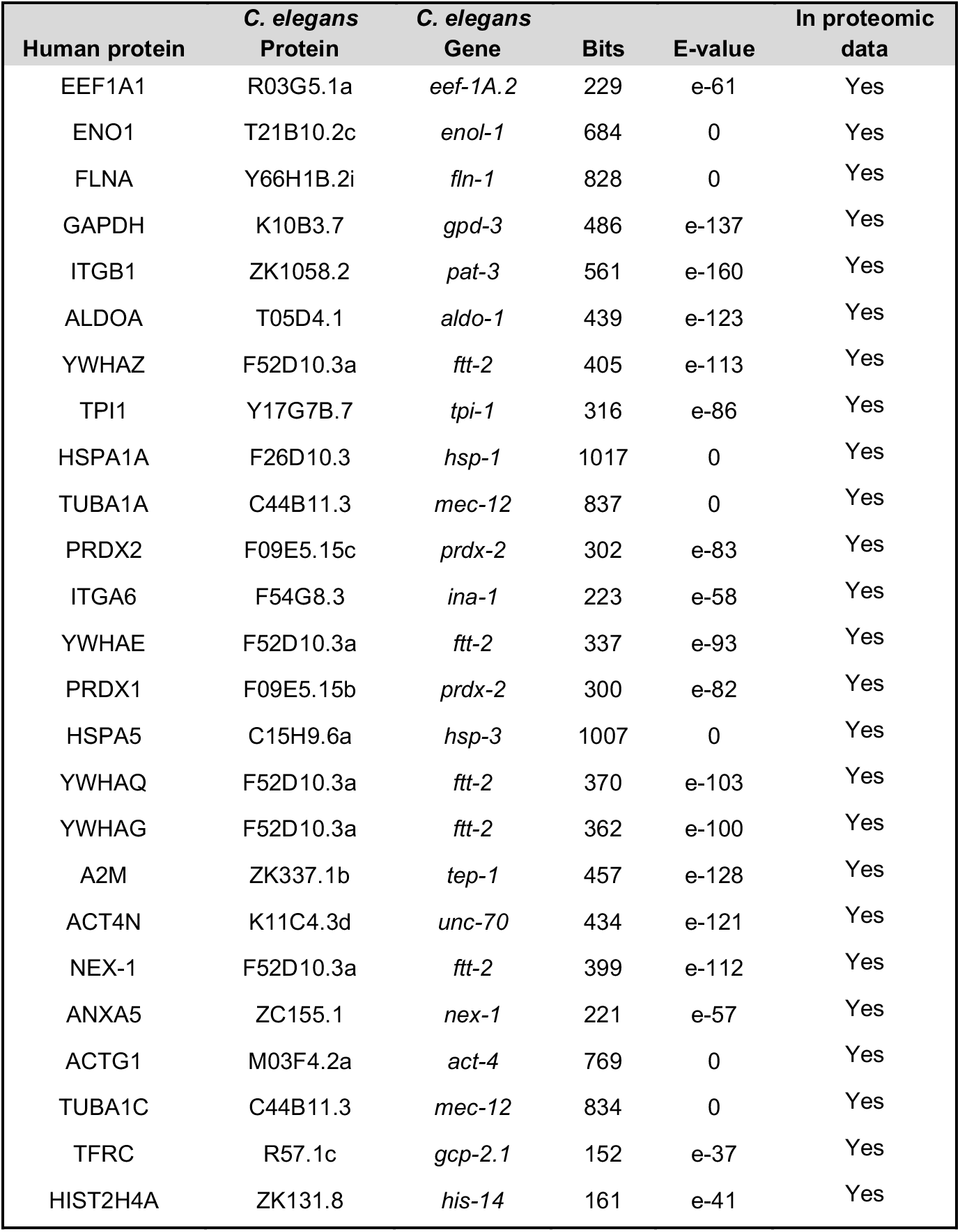
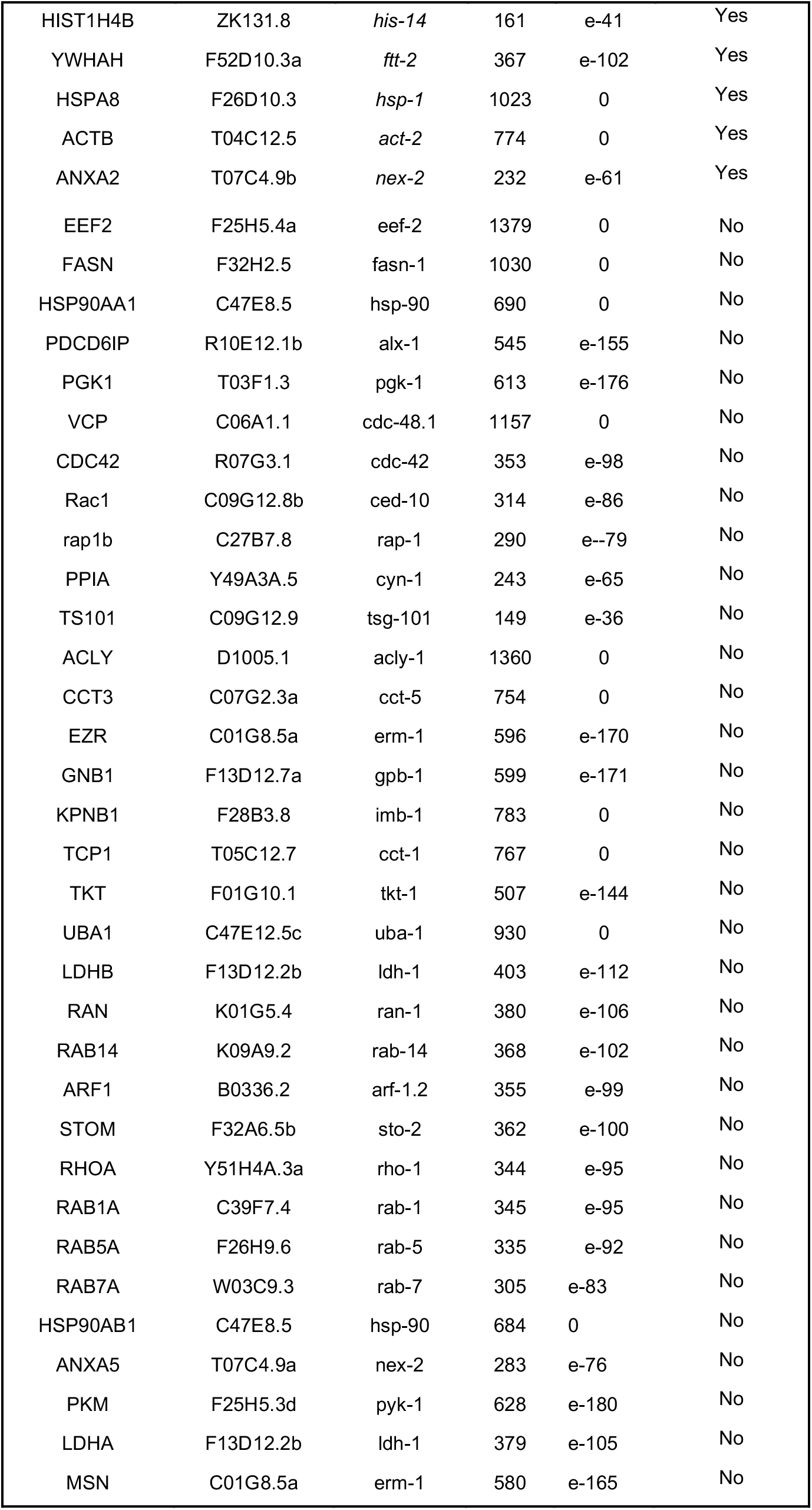
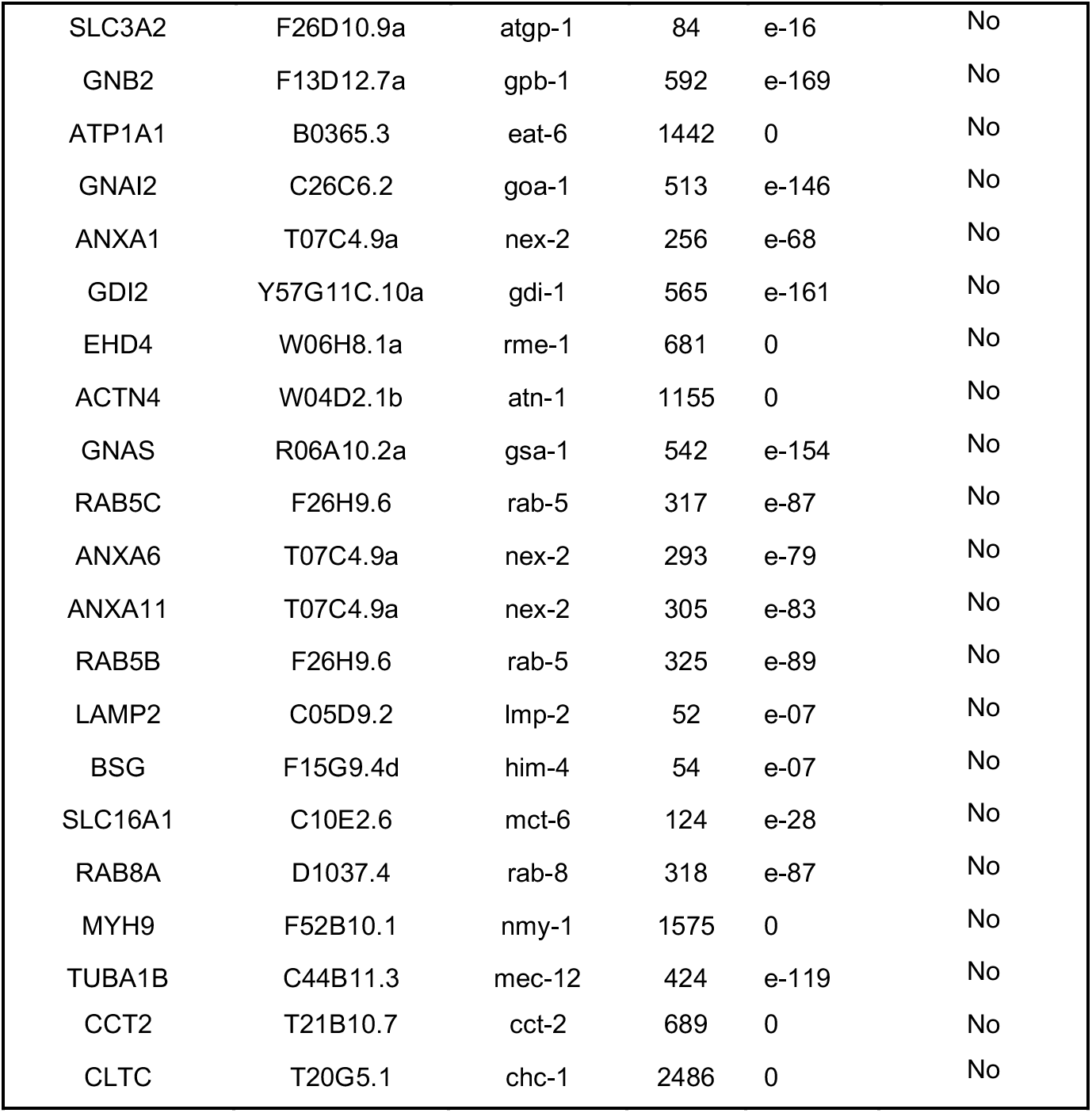
Comparison between proteins identified *in C. elegans* EV proteomics and the top 100 most-identified proteins on Exocarta.org. A) The best reciprocal The *C. elegans* orthologs of the top 100 most-identified human EV protein from Exocarta (exocarta.org) were identified by filtering for proteins pairs that show clear reciprocal best hits. The proteomic data sets were then searched for protein matches. 11 of the 46 human EV proteins (~24%) with best-reciprocal hits were identified in the *C. elegans* EV proteomic datasets. B) Proteomic matches to proteins orthologous to Exocarta top 100. To generate this list *C. elegans* BLAST analysis identified the proteins that showed strongly orthologous peptide sequence (<e-30) to the top 100 most-identified proteins on Exocarta.org. This set of proteins was then compared to the set of proteins identified with unique peptide hits in our EV proteomic experiments. 30 of the 84 human EV proteins that had *C. elegans* proteins with BLAST values < e-30 had matches with proteins identified by unique peptide hits in our proteomics data sets (35%).

